# The Use of Class Imbalanced Learning Methods on ULSAM Data to Predict the Case-Control Status in Genome-Wide Association Studies

**DOI:** 10.1101/2023.01.05.522884

**Authors:** R. Onur Öztornaci, Hamzah Syed, Andrew P. Morris, Bahar Taşdelen

## Abstract

Machine learning (ML) methods for uncovering single nucleotide polymorphisms (SNPs) in genome-wide association study (GWAS) data that can be used to predict disease outcomes are becoming increasingly used in genetic research. Two issues with the use of ML models are finding the correct method for dealing with imbalanced data and data training. This article compares three ML models to identify SNPs that predict type 2 diabetes (T2D) status using the Support vector machine SMOTE (SVM SMOTE), The Adaptive Synthetic Sampling Approach (ADASYN), Random under sampling (RUS) on GWAS data from elderly male participants (165 cases and 951 controls) from the Uppsala Longitudinal Study of Adult Men (ULSAM). It was also applied to SNPs selected by the SMOTE, SVM SMOTE, ADASYN, and RUS clumping method. The analysis was performed using three different ML models: (i) support vector machine (SVM), (ii) multilayer perceptron (MLP) and (iii) random forests (RF). The accuracy of the case-control classification was compared between these three methods. The best classification algorithm was a combination of MLP and SMOTE (97% accuracy). Both RF and SVM achieved good accuracy results of over 90%. Overall, methods used against unbalanced data, all three ML algorithms were found to improve prediction accuracy.

## 1 INTRODUCTION

Genome-wide association studies (GWAS) are the most commonly used strategy for investigating the genetic architecture of common complex diseases. GWAS methodology examines the differences in allele frequencies between individuals affected and unaffected by the disease at single nucleotide polymorphisms (SNPs) across the genome. GWAS data can be used for predicting disease status and classification of cases from controls using machine learning (ML) algorithms. The use of ML algorithms for prediction is increasingly being used over methodology such as polygenic risk scores (Fadista, Manning, Florez, & Groop, 2016), (Szymczak et al., 2009). The application of ML algorithms have some advantages in this situation over classical statistical methodology such as the logistic regression model; i) for the classification of cases and controls, where a large number of possibly correlated variables may be modelled together; ii) a significance threshold is not required; iii) gene-gene interactions can be evaluated and iv) assumptions based on normality and homogeneity of variances are not needed (Cosgun, Limdi, & Duarte, 2011), (Tang, Zhang, Chawla, & Krasser, 2008).

GWAS data are suitable for ML data mining because large amounts of phenotype and genomic data can be studied simultaneously. Large complex GWAS data summarised with data mining approaches can be interpreted quickly and explained by graphics such as decision tree plots. Following the ML procedure, the selection of an appropriate statistical method is necessary to ensure time and cost savings. Methods that use classification are at a disadvantage due to their long calculation time, and increased likelihood of over-fitting and under-fitting. Under-fitting occurs when the model cannot find the intricate patterns hidden in the data. Whereas, when the model learns from large quantities of data, over-fitting is more likely to occur. The likelihood of over-fitting is increased when the dataset is imbalanced (Tang et al., 2008), (Dai, Fu, Zhao, & Zeng, 2021).

Imbalanced data is defined as the category/outcome variable used for classification having unequal classes or differing number of observations. Class imbalance is a problem that arises frequently when applying GWAS for diseases with low prevalence in large cohorts. In this case, it may be a solution to use adjusted the type-I error when using classical statistical methods such as the logistic regression model. However, ML algorithms are not immune to class imbalance. Therefore, datasets need to be balanced with statistically unbiased resampling techniques such as (Synthetic Minority Over-sampling Technique (SMOTE). (Zhou et al., 2018), (Bao et al., 2021).

GWAS typically interrogate millions of SNPs, many of which are in strong linkage disequilibrium (LD) with each other, which poses considerable computational challenges in applying ML methods. LD may affect the classification performance (Meng, Yu, Cupples, Farrer, & Lunetta, 2009). LD-clumping use to remove highly correlated SNPs, may be helpful for increasing classification performance on ML (Thomas et al., 2020). Clumping is a procedure within PLINK that selects the most significant SNPs in LD blocks. By doing this, the data size and the correlation between remaining SNPs is reduced (Purcell et al., 2007). Using clumping with ML is advantageous as it reduces processing time and it is a proven and commonly used procedure for applying ML in GWAS (Kinreich et al., 2021), (K. Y. He, Ge, & He, 2017).

In this paper, ML approaches were applied to a GWAS of type 2 diabetes (T2D) undertaken in the Uppsala Longitudinal Study of Adult Men (ULSAM), which includes 165 cases and 951 controls of European ancestry from Sweden. The objectives of this study are to examine the accuracy, specificity, and sensitivity of three different machine learning approaches and to compare the results. Furthermore, we applied LD clumping in both situations with/without SMOTE and compared the results.

## 2 MATERIAL AND METHODS

When performing GWAS, the entire genome is analysed, to identify SNPs that may predispose a disease of interest. While logistic regression is used for dependent variables in a binary setting, linear regression methods are used for variables in a continuous setting. The datasets used for genetic association studies are high dimensional (Pirooznia et al., 2012). Sample size is very influential for classical statistical approaches like the logistic regression model because a small number of individuals (n) and/or a large number of SNPs (feature, p) can be a problem. The (*X*^T^*X*) parameters w that are used for predicting beta coefficients will not be a singular matrix and the parameters in the regression model cannot be uniquely estimated (Fan & Tang, 2013). However, ML algorithms are less affected by this problem than the classical multivariate approaches because ML algorithms have tuning parameters to optimise and regulate the process. An example of this are penalized likelihood methods (Johnstone & Titterington, 2009), (Nordhausen, 2009). Another issue for both classical and ML models is imbalanced data, where the data collected between groups of samples are different (Draisma et al., 2015). Most often, LD clumping has been used when applying machine learning to GWAS data (Shi, Medway, Brown, Kalsheker, & Morgan, 2011), (Bhowan, Johnston, Zhang, & Yao, 2012). Using clumping has benefits such as reducing calculation time and memory usage. It can be used to eliminate unrelated SNPs that provide redundant information using a defined threshold of LD (usually r^2^) (Bhowan et al., 2012), (Chawla, Bowyer, Hall, & Kegelmeyer, 2002). In this case, the majority class is more likely to be correctly classified than the minority class. In a classification problem, the class that has fewer elements than the other classes are called the minority class. Bias is the main contributor to this classification problem because ML algorithms tend to classify data according to the majority class. SMOTE is a powerful approach for avoiding bias against the minority class (Chawla et al., 2002). We applied ML methods with LD clumping comparing performance with and without SMOTE

### 2.1 SMOTE

Class imbalance occurs in categorical data when one of the classes is less frequent and is a minority class (Japkowicz & Stephen, 2002). This is often a problem in population based GWAS where cases of a disease are much less frequent than controls. This situation poses a problem for machine learning models (Lusa, 2013). High dimensional data sets such as gene expression, GWAS, electronic medical records, and text mining are frequently facing this problem (Turhan, Özkan, Yürekli, Suner, & Doğu, 2020). Synthetic Minority Over-sampling Technique (SMOTE) offers a solution to this problem in conjunction to applying ML. SMOTE increases the number of observations within the minority class, using k-nearest neighbours for balancing the data. SMOTE creates synthetic samples by random interpolation among the nearest neighbours in the same class as the minority class sample(Shrivastava, Jeyanthi, & Singh, 2020). When using interpolation operation, the decision space for the minority class is extended to get the closest possibility to the original variables when creating a new sample in the minority class. The minority class with the new samples have the same expected value as the original minority class samples, however, their variances are not the same (Seo & Kim, 2018) (Hu & Li, 2013), (Z. Zheng, Cai, & Li, 2015).

The new artificial sample *X_new_, X*’ randomly selected neighbor, *X_i_*, for each instance in minority class.

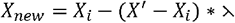

Where λ is a random number in the range [0, 1] and i ∈ [1, *n*], *n* is the number of individuals in the minority class (Z. Zheng et al., 2015).

### 2.2 Support Vector Machine SMOTE

Support vector machine SMOTE (SVM SMOTE) is a technique that based on combination of extrapolation and interpolation that defining boundaries with SVM separation formulas (Q. Wang, Luo, Huang, Feng, & Liu, 2017), N2 (H.-Y. Wang, 2008). Instead of the k-neighbourhood computational interpolation in the SMOTE approach, SVM SMOTE uses support vectors, using interpolation for the majority class, while using extra polarization for the minority class, thus making the sample balanced. Synthetic data will be randomly created along the lines joining each minority class support vector with several its nearest neighbours (Tang et al., 2008).

### 2.3 Adaptive Synthetic Sampling Approach

The Adaptive Synthetic Sampling Approach (ADASYN) is an approach that can be used to solve class imbalance problems based on adaptive minority class generation on data samples. ADASYN increments the minority class by calculating the shuffled and randomly selected majority class distances to form the minority class (H. He, Bai, Garcia, & Li, 2008) [N4]. While increasing the number of minority class elements, it not only reduces bias by randomizing the data instead of choosing the distances of those close to the majority class, but also increases the number of minority class elements by considering the majority class elements that are difficult to include in the calculations (Alhudhaif, 2021).

### 2.4. Random Under Sampling

Random under sampling (RUS) technique is a method used to provide balance between minority class and majority class in large sample size data with class imbalance. In this method, instead of increasing the number of minority class elements, resampling is done by randomly selecting the majority class elements to equal the number of minority class elements (Zuech, Hancock, & Khoshgoftaar, 2021). While applying this method, no mathematical approach is applied, and majority class elements are chosen randomly. Therefore, in this case, data with potentially important information may be deleted (Razavi-Far, Farajzadeh-Zanajni, Wang, Saif, & Chakrabarti, 2019).

### 2.5. Machine Learning General Concepts

Machine learning is a rapidly evolving discipline that involves the examination, learning and development of algorithms to improve computational performance when analysing data. These algorithms help us make the most accurate hypothesis-driven decisions. The two main categories of machine learning are supervised and unsupervised learning (Han, Pei, & Kamber, 2011). When applying ML methods, data should be divided into training and testing sets. Training is a data subset to train the algorithm, and the test data is for evaluating the performance of the algorithm. Test sets must be independent of the training set to avoid over-fitting. Test sets cannot be used for training the algorithm (Alpaydin, 2020).

Supervised learning is generally concerned with examining classification problems. We can think of classification models in two steps: the first step is to classify the data according to the existing characteristics, constraints, and conditions, and the second step is to test the accuracy of the classes and the validity of the model (Han et al., 2011). If we want to define the classification, simplistically, we can say that it is the process of predicting the classes of data. It is the process of segmenting similar objects, observations, and events according to a specific purpose (Alpaydin, 2020).

### 2.3 Random Forests

Random forests (RF), which were introduced by Breiman *et al*. (Breiman, 2001) in 2001, are combinations of several random decision trees, with each tree sampled independently and with the same distribution. The main objective is to achieve higher accuracy for prediction. All trees are constructed with a different bootstrap sample selection from the original data set (Chen & Ishwaran, 2012). Random forests are generated using the Classification and Regression Tree (CART) algorithm, and an information gain criterion is used for splitting each node. Although RF can be used for data with many variables, it requires a lot of available computational memory (M. Pal, 2005). The Gini index is used for building the sub-trees within the random forest. The Gini index formulation can be written as the following, where T represents the training set and (*f*(*G_i_.T*)|*T*|) is the probability of the cases selected belonging to the class *G_i_* (Strobl & Zeileis, 2008).

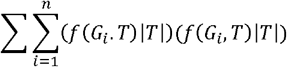

### 2.4 Support Vector Machine

Support vector machines (SVM) were developed by Vapnik *et al*. (Guyon, Weston, Barnhill, & Vapnik, 2002). SVM algorithms can be used to find the solution of both clustering (unsupervised) and classification (supervised) problems. Consider drawing a border separating two groups along a plane. SVM determines how this border is drawn (Furey et al., 2000; Mammone, Turchi, & Cristianini, 2009). SVM algorithms search for the best solution to the classification problem by utilising optimisation principles which are useful for “big data” evaluation (Nitze, Schulthess, & Asche, 2012), (Mieth et al., 2016). SVMs can be used for classification, dimension reduction, and SNP selection, by applying one of three different kernels: linear kernel, polynomial kernel, or Radial basis function kernel (RBF)/Gaussian Kernel. RBF is used when solving non-linear problems (Ng & Mishra, 2007), (Statnikov, Aliferis, Tsamardinos, Hardin, & Levy, 2005), (Deng, Shen, Wang, & Zhang, 2020). It is crucial to draw a small margin amongst the two classes to reduce the number of errors produced through classification. If we select two points for the realisation of linear separation and define them as k and l, we maximise the equations *wx_k_*, + *b* = – 1 *and wx_l_* + *b* = +1 through subtraction.

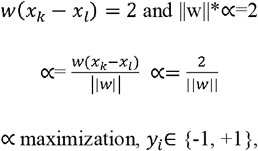

Conditionally, if *y_i_*=+1 then *w^T^x* — *b* ≥ 1 or if *y_i_*=-1 then *w^T^x* – *b* ≤ −1. From this, we can deduce that the minimum is calculated using, 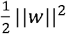 Where, w is a weight vector, b is bias, x is the input vector, and *y_i_* is the I’th target (i.e. in this case, 1 or −1).

### 2.5 Multi-Layer Perceptron

Multilayer perceptron (MLP) is a method that can be used for both classification and clustering. It is a method inspired by the human brain (S. K. Pal & Mitra, 1992). The MLP follows a neural network structure which consists of three different layers; i) input, ii) hidden (one or more), and iii) output. MLP, is a feed-forward method (Fergus et al., 2018), where hidden layers provide a transmission from the input layer to the output layer to classify variables. MLP offers a solution for non-linear classification problems, because it uses the delta learning rule. This rule is based on sigmoid rules such as a logistic function (Elmas & Uygulamaları, 2007), (Pedregosa et al., 2011). The learning function of MLP is represented as: the weights. Delta rules are defined by:

W and are model parameters, G represents the Genotypes and W represents the weights. Delta rules are defined by:

Where, is the learning rate, is the derivative of the target output. is the weighted sum of the neuron’s inputs and is the actual output defined as.

### 2.6 Logistic Regression

Logistic regression (LR) is a statistical method used in binary data to determine the relationship between SNPs and disease in GWAS. Since LR is the classical statistical approach, it is used to measure the p-value and odds ratio ratios and the relationship between SNPs and disease (Staley et al., 2017). Generally, SNPs associated with the disease are determined with the aid of a Manhattan plot, along the threshold of 10^-8^. Since LR is a method used as a classification method, in this study, machine learning methods and classification success were compared (Wakefield, 2009).

### 2.7 Performance Measurement Metrics

We have assessed the prediction of these ML algorithms using five complementary metrics. Table 1 below defines the confusion matrix terms.

**Table 1.** Confusion matrix definitions for the predicted and real class.

| Confusion Matrix | Predicted Class |  |  |
| --- | --- | --- | --- |
|  | Status | Control | Case |
| Real Class | Control | True Negative (TN) | False Positive (FP) |
|  | Case | False Negative (FN) | True Positive (TP) |

Accuracy is determined by dividing all data by the predicted values. This method, however, is not a sufficient measurement when using imbalanced data.

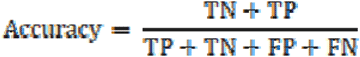

Sensitivity is calculated by dividing the true positive cases by all predicted positive cases.

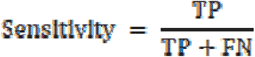

Specificity is calculated by dividing the true negative cases by all predicted negative cases.

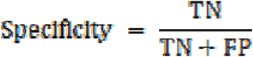

The Positive Predictive Value is the percentage of cases that are true positives as predicted by the ML algorithms.

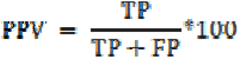

The Negative Predictive Value is the percentage of controls that are true controls who are predicted by the ML algorithms.

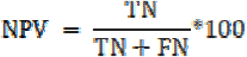

The F1 score (also known as an F-score or F-measure) is a measure of accuracy. When calculating the F1 score, both precision and recall metrics are used.

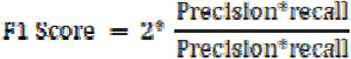

The F1 score is used for the evaluation of data that has class imbalance. This metric is widely used for the evaluation of ML models (Chicco & Jurman, 2020), (Korkmaz, 2020).

### 2.8 ULSAM Study

The Uppsala Longitudinal Study of Adult Men (ULSAM) is an investigation of healthy elderly men in the Uppsala region of Sweden (Lithell et al., 2000). It was initiated as a health screen to identify metabolic risk factors for cardiovascular disease. In 1970, all 50-year-old men living in Uppsala were invited to participate, of whom 82% initially agreed to participate, and were subsequently invited back for further study at ages 60, 70 and 77. At each visit, a wide range of phenotypes were collected, including blood pressure, insulin metabolism, weight and height, lipid markers, diet, cognitive function, and socio-economic factors. At age 70, participants were given a glucose tolerance test and insulin clamp to measure insulin resistance. T2D case status was defined by doctor-diagnosed disease or fasting whole blood glucose >6.1mmol/l, with all non-cases defined as controls. A total of 1178 participants were genotyped with the Illumina 2.5M Omni array and Illumina CardioMetaochip (Lithell et al., 2000). Samples were excluded if the call rate was less than 95%, if they had extreme heterozygosity (>3 SD from the mean), if they were of non-European ancestry, or if they were female on the basis of X chromosome data. SNP quality control measures included exact p-value for deviation from (Hardy Weinberg Equilibrium) HWE <10^-6^, call rate less than 95% (or less than 99% for SNPs with (Minor Allele Frequency) MAF <5%), and MAF <1%. Multidimensional scaling (MDS) of a genetic relatedness matrix from LD-pruned autosomal data was performed to obtain principal components to adjust for population structure.

## 3 RESULTS

### 3.1 Machine Learning Results without Clumping

The ULSAM GWAS data were divided into two parts; 70% training set and 30% testing set in preparation for the supervised learning algorithms for classifying cases and controls. 10-fold cross-validation was performed to avoid over-fitting (Han et al., 2011). A total of 399,935 SNPs with no missing genotype information for each of the 1116 samples (165 cases and 951 controls) were used in the analysis. After using SMOTE for the adjustment of imbalanced classes, there were 1902 individuals, equally divided between the two classes (951 cases and 951 controls. After using SVM SMOTE for the adjustment of imbalanced classes, there were individuals, 1552 equally divided between the two classes (776 cases and 776 controls). After using ADASYN for the adjustment of imbalanced classes, there were individuals, 1872 equally divided between the two classes (936 cases and 936 controls). After using RUS for the adjustment of imbalanced classes, there were individuals, 466 equally divided between the two classes (233 cases and 233 controls). The choice of tuning parameters affects both the sensitivity and classification performance independently (Lavesson & Davidsson, 2006). Therefore, we used the optimal tuning parameters according to the instructions in the Scikit-learn package documentation for Python 3.7 (Pedregosa et al., 2011), (Van Rossum, 2007).

When comparing the methods used to eliminate the whole class imbalance and the original data, SMOTE method gives the best results for SVM with the highest accuracy rate (91%) and F1 score (90%). The SMOTE method is followed by the ADASYN method with an accuracy of 90% and an F1 score of 89%. The RUS method, on the other hand, had the worst results.

When comparing the methods used to eliminate the whole class imbalance and the original data, SMOTE method gives the best results for support vector machines with the highest accuracy rate (91%) and F1 score (90%). The SMOTE method is followed by the ADASYN method with an accuracy of 90% and an F1 score of 89%. The RUS method, on the other hand, had the worst results. (Table 2).

**Table 2.** The Performances of Support Vector Machine with Imbalanced Learning Methods.

| *Imbalanced Learning Methods* |  | *Prediction Class* |  | *PPV\** | *NPV\*\** | *Sensitivity* | *Specificity* | *F1 Score* | *Accuracy* |
| --- | --- | --- | --- | --- | --- | --- | --- | --- | --- |
| *Number of SNP 399935* |  |  |  |  |  |  |  |  |  |
| *SVM* |  | *Controls* | *Cases* |  |  |  |  |  |  |
| **Reel Class** | *Controls* | 278 | 0 | 0.00 | 0.82 | 0.00 | 1.00 | 0.00 | 0.82 |
|  | *Cases* | 57 | 0 |  |  |  |  |  |  |
| *SMOTE* |  | *Controls* | *Cases* |  |  |  |  |  |  |
| **Reel Class** | *Controls* | 287 | 0 | 1.00 | 0.85 | 0.83 | 1.00 | 0.90 | 0.91 |
|  | *Cases* | 47 | 237 |  |  |  |  |  |  |
| *SVM SMOTE* |  | *Controls* | *Cases* |  |  |  |  |  |  |
| **Reel Class** | *Controls* | 293 | 0 | 1.00 | 0.83 | 0.67 | 1.00 | 0.80 | 0.87 |
|  | *Cases* | 56 | 117 |  |  |  |  |  |  |
| *ADASYN* |  | *Controls* | *Cases* |  |  |  |  |  |  |
| **Reel Class** | *Controls* | 285 | 0 | 1.00 | 0.84 | 0.81 | 1.00 | 0.89 | 0.90 |
|  | *Cases* | 51 | 226 |  |  |  |  |  |  |
| *RUS* |  | *Controls* | *Cases* |  |  |  |  |  |  |
| **Reel Class** | *Controls* | 47 | 0 | ** | 0.47 | 0.00 | 1.00 | 0.00 | 0.47 |
|  | *Cases* | 53 | 0 |  |  |  |  |  |  |

\* PPV: Positive Predictive Value, \*\* NPV: Negative Predictive Value

As can be seen with the roc curve, it is seen that the classification success of the results of the SVM, which is used with the RUS method without any correction, is low (Fig.1).

**Figure 1.**
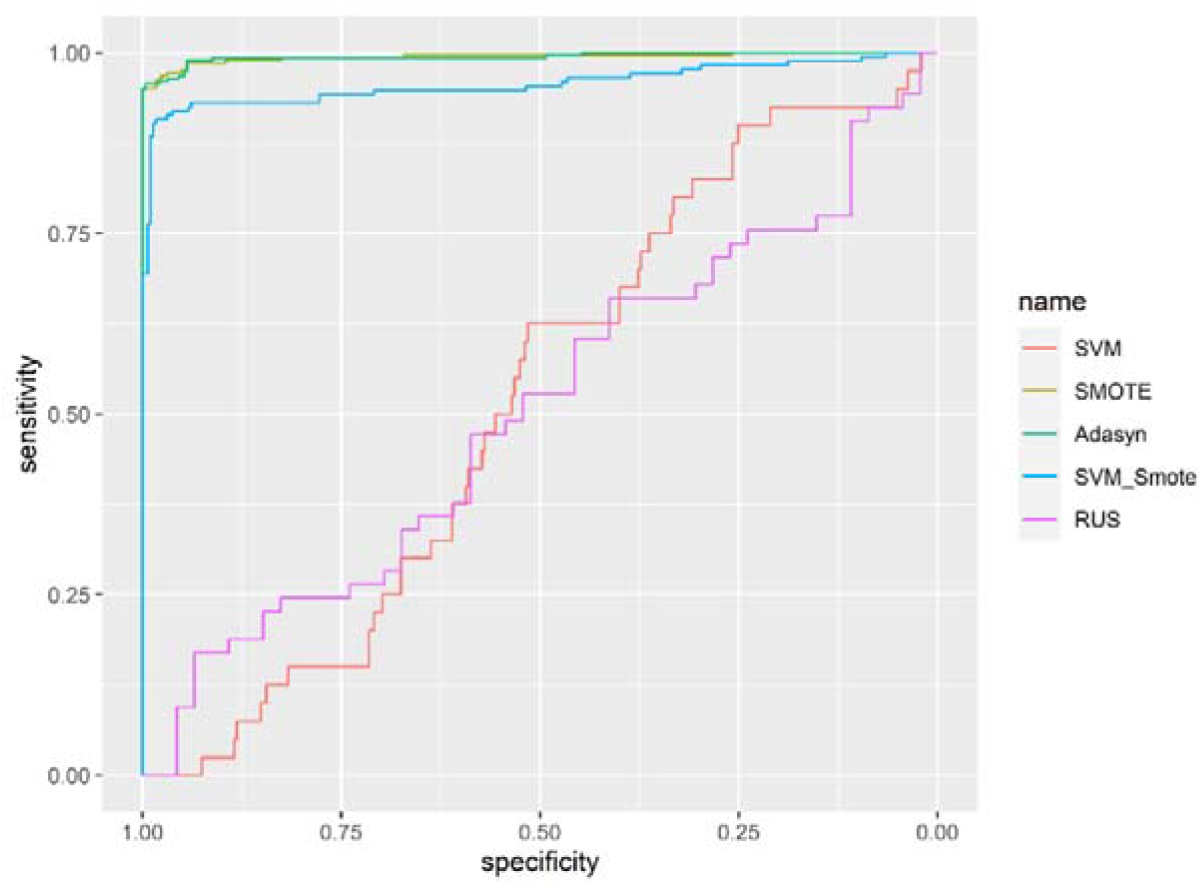
ROC Curve: Comparison of the performances of Support Vector Machine with Imbalanced Learning Methods

When comparing the methods used to eliminate the whole class imbalance and the original data, the SMOTE method and ADASYN are the methods that give the best results for the random forest with the highest accuracy rates (92%) and approximately F1 scores (92%). Therefore, both methods can be used interchangeably. Although the SVM SMOTE method achieved accuracy with a 5% difference compared to the method whose class Imbalance problem was not resolved, it achieved better results in terms of F1 score and sensitivity and specificity than the case without class imbalance. Although the RUS method has the lowest accuracy rate, it has a sensitivity difference of 35% compared to the original data (Table 3).

**Table 3.** The Performances of Random Forest with Imbalanced Learning Methods.

| *Imbalanced Learning Methods*<br>*Number of SNP 399935* |  | *Prediction Class* |  | *PPV\** | *NPV\*\** | *Sensitivity* | *Specificity* | *F1 Score* | *Accuracy* |
| --- | --- | --- | --- | --- | --- | --- | --- | --- | --- |
| **RF** |  | *Controls* | *Cases* |  |  |  |  |  |  |
| **Reel Class** | *Controls* | 278 | 0 | 0.00 | 0.82 | 0.00 | 1.00 | 0.00 | 0.82 |
|  | *Cases* | 57 | 0 |  |  |  |  |  |  |
| **SMOTE** |  | *Controls* | *Cases* |  |  |  |  |  |  |
| **Reel Class** | *Controls* | 286 | 1 | 0.99 | 0.87 | 0.85 | 0.99 | 0.92 | 0.92 |
|  | *Cases* | 40 | 244 |  |  |  |  |  |  |
| **SVM SMOTE** |  | *Controls* | *Cases* |  |  |  |  |  |  |
| **Reel Class** | *Controls* | 290 | 3 | 0.97 | 0.84 | 0.68 | 0.98 | 0.80 | 0.87 |
|  | *Cases* | 54 | 119 |  |  |  |  |  |  |
| **ADASYN** |  | *Controls* | *Cases* |  |  |  |  |  |  |
| **Reel Class** | *Controls* | 285 | 0 | 1.00 | 0.86 | 0.84 | 1.00 | 0.91 | 0.92 |
|  | *Cases* | 43 | 234 |  |  |  |  |  |  |
| **RUS** |  | *Controls* | *Cases* |  |  |  |  |  |  |
| **Reel Class** | *Controls* | 26 | 21 | 0.47 | 0.43 | 0.35 | 0.55 | 0.40 | 0.45 |
|  | *Cases* | 34 | 19 |  |  |  |  |  |  |

\* PPV: Positive Predictive Value, \*\* NPV: Negative Predictive Value

As can be seen from the Roc curve, it is seen that the classification success of the RF results used without any correction with the RUS method is low, and at the same time, the lines for the SMOTE and ADASYN methods intersect at almost the same point, therefore, the ADASYN and SMOTE methods are almost equal in terms of all metrics (Fig. 2).

**Figure 2.**
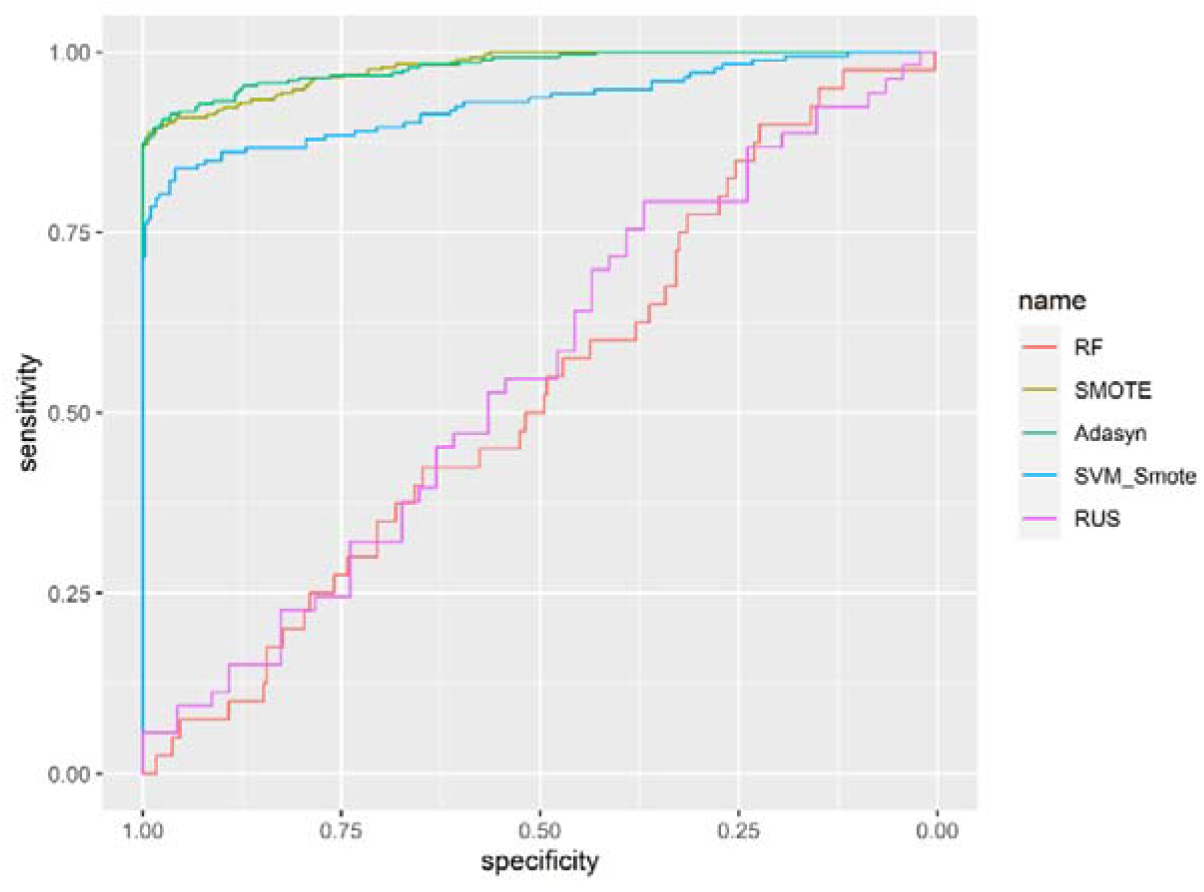
ROC Curve: Comparison of the performances of Random Forest with Imbalanced Learning Methods

Among the methods used to correct class imbalance for MLP, the SMOTE method has the best result with the highest accuracy rate (97%) and F1 score (97%). The SVM SMOTE method, on the other hand, follows the SMOTE method with 92% accuracy and 88% F1 score rate. Although the ADASYN method is not as high in accuracy as SMOTE and SVM SMOTE, it has a very good F1 score (93%). The RUS method, on the other hand, was seen as a bad method (47%) with accuracy and (7%) F1 score (Table 4).

**Table 4.** The Performances of Multi-layer Perceptron with Imbalanced Learning Methods.

| *Imbalanced Learning Methods*<br>*Number of SNP 399935* |  | *Prediction Class* |  | *PPV\** | *NPV\*\** | *Sensitivity* | *Specificity* | *F1 Score* | *Accuracy* |
| --- | --- | --- | --- | --- | --- | --- | --- | --- | --- |
| **MLP** |  | *Controls* | *Cases* |  |  |  |  |  |  |
| **Reel Class** | *Controls* | 277 | 1 | 0.00 | 0.82 | 0.00 | 0.99 | 0.00 | 0.82 |
|  | *Cases* | 57 | 0 |  |  |  |  |  |  |
| **SMOTE** |  | *Controls* | *Cases* |  |  |  |  |  |  |
| **Reel Class** | *Controls* | 282 | 5 | 0.98 | 0.96 | 0.96 | 0.98 | 0.97 | 0.97 |
|  | *Cases* | 9 | 275 |  |  |  |  |  |  |
| **SVM SMOTE** |  | *Controls* | *Cases* |  |  |  |  |  |  |
| **Reel Class** | *Controls* | 290 | 3 | 0.97 | 0.89 | 0.80 | 0.98 | 0.88 | 0.92 |
|  | *Cases* | 33 | 140 |  |  |  |  |  |  |
| **ADASYN** |  | *Controls* | *Cases* |  |  |  |  |  |  |
| **Reel Class** | *Controls* | 285 | 0 | 1.00 | 0.89 | 0.88 | 1.00 | 0.93 | 0.84 |
|  | *Cases* | 32 | 245 |  |  |  |  |  |  |
| **RUS** |  | *Controls* | *Cases* |  |  |  |  |  |  |
| **Reel Class** | *Controls* | 45 | 2 | 0.50 | 0.46 | 0.03 | 0.95 | 0.07 | 0.47 |
|  | *Cases* | 51 | 2 |  |  |  |  |  |  |

\* PPV: Positive Predictive Value, \*\* NPV: Negative Predictive Value

As can be seen from the Roc curve, SMOTE, SVM SMOTE, and ADASYN methods have very high classification success for MLP. The original data and RUS methods showed poor classification performance (fig.3.).

**Figure 3.**
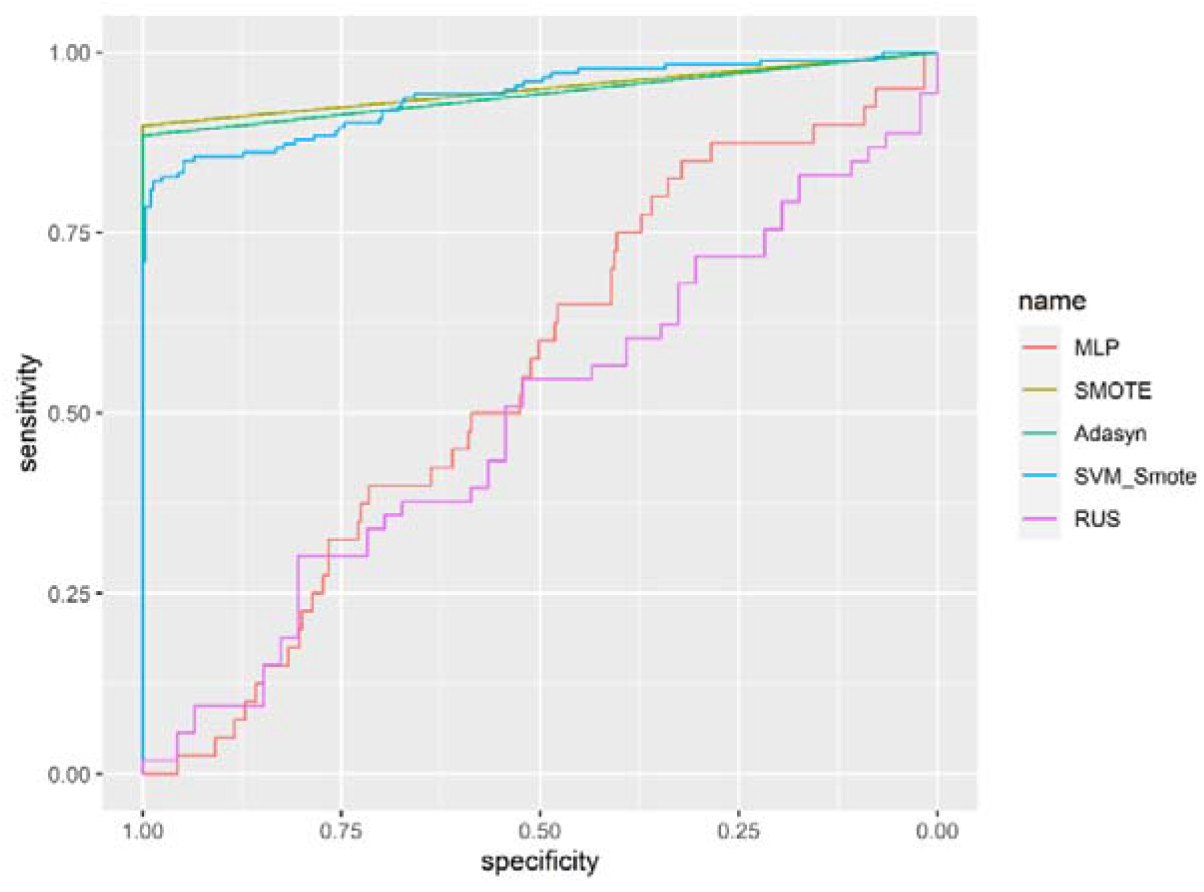
ROC Curve: Comparison of the performances of Multi-Layer Perceptron with Imbalanced Learning Methods

The logistic regression results applied to the data with class imbalance were found to be quite similar to all machine learning results, with an accuracy rate of 82% and an F1 score of 0.00. The ADASYN method achieved very close results to the SMOTE method, with an accuracy of 95% and an F1 score of 93%. The SVM SMOTE method can be considered as an alternative to these two methods with 90% accuracy and 84% F1 score. Although the RUS method, like all other machine learning methods, had poor results, it achieved a 41% higher sensitivity rate than the data with unbalanced classes (Table 5).

**Table 5.** The Performances of Logistic Regression with Imbalanced Learning Methods.

| Imbalanced Learning Methods |  | Prediction Class |  | PPV* | NPV** | Sensitivity | Specificity | F1 Score | Accuracy |
| --- | --- | --- | --- | --- | --- | --- | --- | --- | --- |
| Number of SNP 399935 |  |  |  |  |  |  |  |  |  |
| LR |  | Controls | Cases |  |  |  |  |  |  |
| Reel Class | Controls | 276 | 2 | 0.00 | 0.82 | 0.00 | 0.99 | 0.00 | 0.82 |
|  | Cases | 57 | 0 |  |  |  |  |  |  |
| SMOTE |  | Controls | Cases |  |  |  |  |  |  |
| Reel Class | Controls | 284 | 0 | 0.90 | 1.00 | 1.00 | 0.91 | 0.95 | 0.95 |
|  | Cases | 26 | 262 |  |  |  |  |  |  |
| SVM SMOTE |  | Controls | Cases |  |  |  |  |  |  |
| Reel Class | Controls | 291 | 2 | 0.98 | 0.86 | 0.74 | 0.99 | 0.84 | 0.90 |
|  | Cases | 44 | 129 |  |  |  |  |  |  |
| ADASYN |  | Controls | Cases |  |  |  |  |  |  |
| Reel Class | Controls | 285 | 0 | 1.00 | 0.89 | 0.87 | 1.00 | 0.93 | 0.93 |
|  | Cases | 35 | 242 |  |  |  |  |  |  |
| RUS |  | Controls | Cases |  |  |  |  |  |  |
| Reel Class | Controls | 28 | 19 | 0.53 | 0.47 | 0.41 | 0.59 | 0.46 | 0.50 |
|  | Cases | 31 | 22 |  |  |  |  |  |  |

\* PPV: Positive Predictive Value, \*\* NPV: Negative Predictive Value

For LR, the SMOTE, ADASYN, and SVM SMOTE methods, which are used to eliminate class imbalance, yield almost identical metrics, resulting in very similar results for the entire Roc curve. For RUS and the original data, the ROC curve clearly showed that both methods failed in classification (fig. 4.).

**Figure 4.**
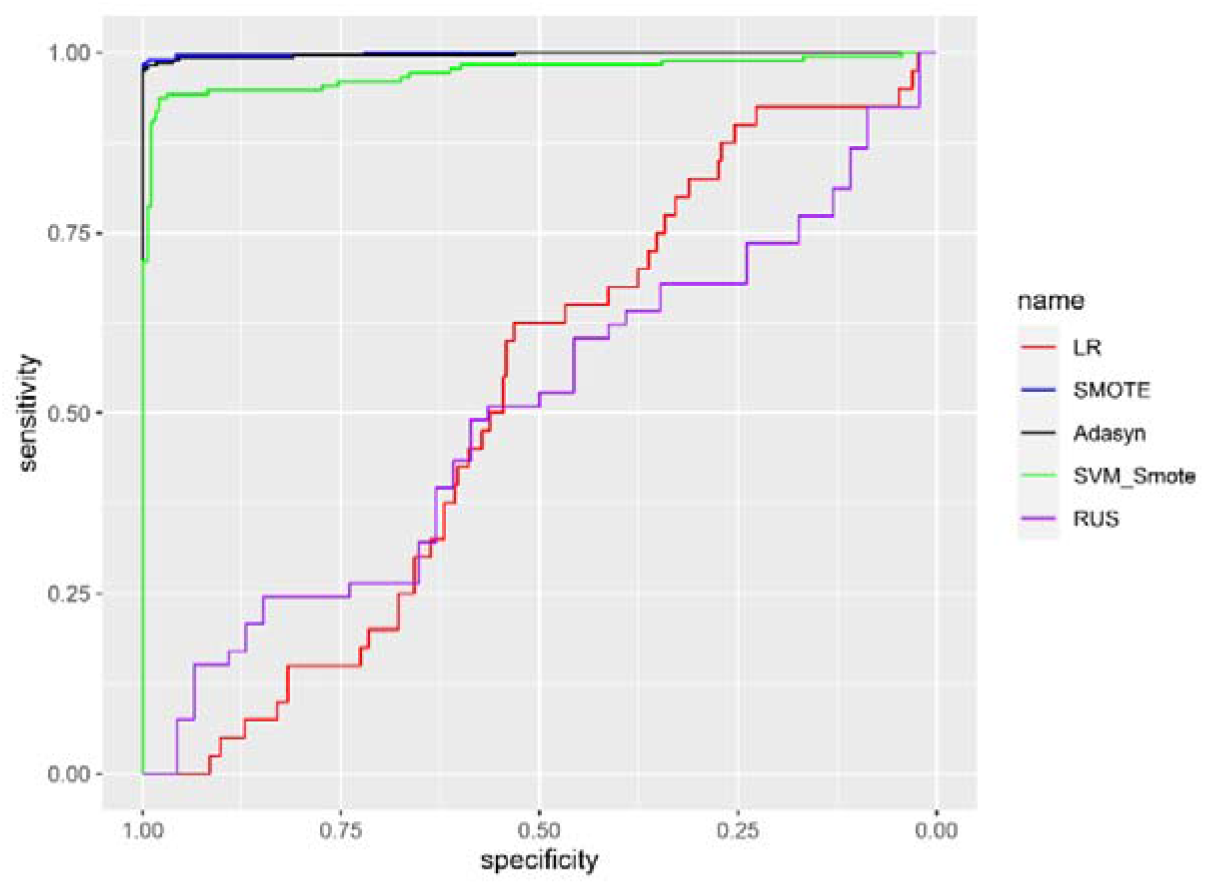
ROC Curve: Comparison of the Performances of Logistic Regression with Imbalanced Learning Methods

### 3.2 Machine Learning Results with Clumping

Following the clumping procedure in PLINK (Purcell et al., 2007), the thresholds were chosen as 0.0001, 0.01, 0.50 and 250 for the parameters p1, p2, r2 and kb, respectively. p1: Significance threshold for index SNPs, p2: Secondary significance threshold for clumped SNPs, r2: LD threshold for clumping, kb: Physical distance threshold for clumping. PLINK (Purcell et al., 2007) was used to LD clump SNPs, using an r2 threshold of 0.50 in windows of 250kb, based on a significance threshold of p<0.01 for index SNPs and p<0.0001 for clumped SNPs. In total, 29 SNPs with very low correlation with one another were obtained.

Data with class imbalances are similar to results before using clumping (e.g., 83% accuracy, 0% F1 score). All the Class Imbalance resolved methods significantly increased the sensitivity measure. The lowest sensitivity rate was found with RUS (72%). All imbalanced learning methods significantly increased the F1 score. SMOTE has the highest F1 score with 82%. (Table 6).

**Table 6.** The Performances of Support Vector Machine with Imbalanced Learning Methods with using Clumped SNPs.

| *Imbalanced Learning Methods* |  | *Prediction Class* |  | *PPV\** | *NPV\*\** | *Sensitivity* | *Specificity* | *F1 Score* | *Accuracy* |
| --- | --- | --- | --- | --- | --- | --- | --- | --- | --- |
| *Number of SNP 29* |  |  |  |  |  |  |  |  |  |
| *SVM* |  | *Controls* | *Cases* |  |  |  |  |  |  |
| **Reel Class** | *Controls* | 278 | 0 | 0.00 | 0.83 | 0.00 | 1.00 | 0.00 | 0.83 |
|  | *Cases* | 57 | 0 |  |  |  |  |  |  |
| *SMOTE* |  | *Controls* | *Cases* |  |  |  |  |  |  |
| **Reel Class** | *Controls* | 190 | 84 | 0.76 | 0.85 | 0.89 | 0.69 | 0.82 | 0.79 |
|  | *Cases* | 34 | 263 |  |  |  |  |  |  |
| *SVM SMOTE* |  | *Controls* | *Cases* |  |  |  |  |  |  |
| **Reel Class** | *Controls* | 225 | 53 | 0.81 | 0.77 | 0.77 | 0.81 | 0.79 | 0.79 |
|  | *Cases* | 67 | 226 |  |  |  |  |  |  |
| *ADASYN* |  | *Controls* | *Cases* |  |  |  |  |  |  |
| **Reel Class** | *Controls* | 201 | 83 | 0.75 | 0.83 | 0.86 | 0.71 | 0.80 | 0.78 |
|  | *Cases* | 42 | 249 |  |  |  |  |  |  |
| *RUS* |  | *Controls* | *Cases* |  |  |  |  |  |  |
| **Reel Class** | *Controls* | 44 | 12 | 0.72 | 0.79 | 0.72 | 0.79 | 0.72 | 0.76 |
|  | *Cases* | 12 | 31 |  |  |  |  |  |  |

\* PPV: Positive Predictive Value, \*\* NPV: Negative Predictive Value

In terms of the Clumping and SVM method, the classification performances of all the methods used to eliminate the class imbalance are very close to each other. Because, when metric values such as sensitive and specificity are examined, it is seen that the results are close to each other (fig. 5).

**Figure 5.**
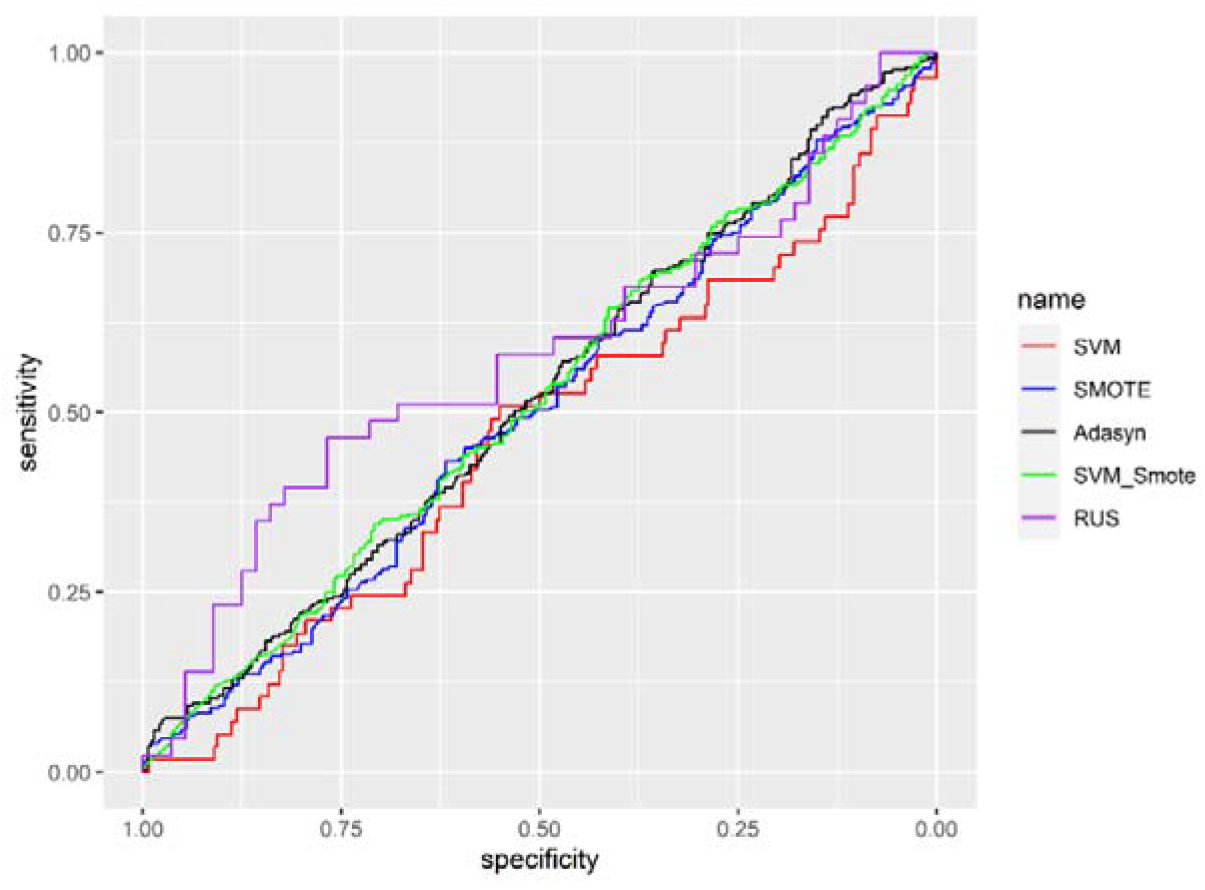
ROC Curve: Comparison of the performances of Support Vector Machine with Imbalanced Learning Methods with using Clumped SNPs

The RF used with the SMOTE method achieved the best result in terms of F1 score (82%). There is no difference between the clumping method and the use of all SNPs in data with class imbalance, as in SVM. A good PPV value was obtained with SVM SMOTE (90%). The ADASYN method and the RUS methods obtained very close results with an accuracy rate of 75% (Table 7).

**Table 7.** The Performances of Random Forest with Imbalanced Learning Methods with using Clumped SNPs.

| *Imbalanced Learning Methods* |  | *Prediction Class* |  | *PPV\** | *NPV\*\** | *Sensitivity* | *Specificity* | *F1 Score* | *Accuracy* |
| --- | --- | --- | --- | --- | --- | --- | --- | --- | --- |
| *Number of SNP 29* |  |  |  |  |  |  |  |  |  |
| **RF** |  | *Controls* | *Cases* |  |  |  |  |  |  |
| **Reel Class** | *Controls* | 278 | 0 | 0.00 | 0.82 | 0.00 | 1.00 | 0.00 | 0.82 |
|  | *Cases* | 57 | 0 |  |  |  |  |  |  |
| **SMOTE** |  | *Controls* | *Cases* |  |  |  |  |  |  |
| **Reel Class** | *Controls* | 184 | 90 | 0.75 | 0.84 | 0.89 | 0.67 | 0.81 | 0.78 |
|  | *Cases* | 34 | 263 |  |  |  |  |  |  |
| **SVM SMOTE** |  |  |  |  |  |  |  |  |  |
| **Reel Class** | *Controls* | 260 | 18 | 0.90 | 0.65 | 0.53 | 0.94 | 0.67 | 0.73 |
|  | *Cases* | 138 | 155 |  |  |  |  |  |  |
| **ADASYN** |  | *Controls* | *Cases* |  |  |  |  |  |  |
| **Reel Class** | *Controls* | 216 | 68 | 0.76 | 0.74 | 0.74 | 0.76 | 0.75 | 0.75 |
|  | *Cases* | 76 | 215 |  |  |  |  |  |  |
| **RUS** |  | *Controls* | *Cases* |  |  |  |  |  |  |
| **Reel Class** | *Controls* | 39 | 17 | 0.67 | 0.83 | 0.81 | 0.70 | 0.74 | 0.75 |
|  | *Cases* | 8 | 35 |  |  |  |  |  |  |

\* PPV Positive Predictive Value, \*\* NPV: Negative Predictive Value

It is seen in the graph that the results very close to the use of the Clumping SVM method are also obtained with the RF method (fig. 6).

**Figure 6.**
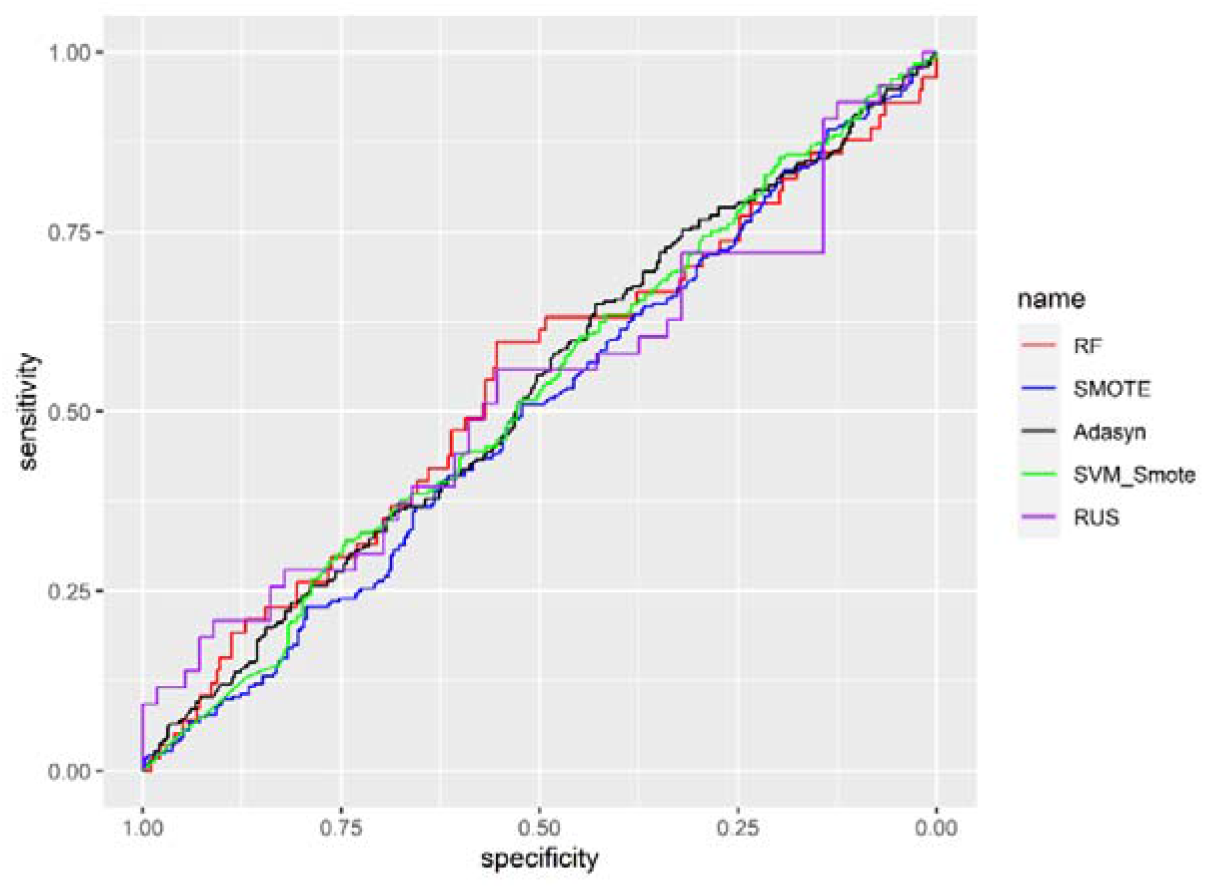
ROC Curve: Comparison of the performances of Random Forest with Imbalanced Learning Methods with using Clumped SNPs

Compared to SVM and RF, the use of clumping with MLP gave relatively better results with an accuracy rate of 82% and an F1 score of 43%. However, when the class imbalance problem was resolved, F1 scores above 80% were obtained with SMOTE, SVM SMOTE and ADASYN. It should be considered that the RUS method also achieved an F1 score of 63%, although it had a low accuracy rate of 69% compared to the case where the class imbalance was not resolved (Table 8).

**Table 8.** The Performances of Multi-Layer Perceptron with Imbalanced Learning Methods with using Clumped SNPs.

| *Imbalanced Learning Methods*<br>*Number of SNP 29* |  | *Prediction Class* |  | *PPV\** | *NPV\*\** | *Sensitivity* | *Specificity* | *F1 Score* | *Accuracy* |
| --- | --- | --- | --- | --- | --- | --- | --- | --- | --- |
| **MLP** |  | *Controls* | *Cases* |  |  |  |  |  |  |
| **Reel Class** | *Controls* | 251 | 27 | 0.46 | 0.88 | 0.40 | 0.90 | 0.43 | 0.82 |
|  | *Cases* | 34 | 23 |  |  |  |  |  |  |
| **SMOTE** |  | *Controls* | *Cases* |  |  |  |  |  |  |
| **Reel Class** | *Controls* | 210 | 64 | 0.79 | 0.80 | 0.82 | 0.77 | 0.81 | 0.80 |
|  | *Cases* | 52 | 245 |  |  |  |  |  |  |
| **SVM SMOTE** |  | *Controls* | *Cases* |  |  |  |  |  |  |
| **Reel Class** | *Controls* | 221 | 57 | 0.81 | 0.81 | 0.82 | 0.79 | 0.81 | 0.81 |
|  | *Cases* | 53 | 240 |  |  |  |  |  |  |
| **ADASYN** |  | *Controls* | *Cases* |  |  |  |  |  |  |
| **Reel Class** | *Controls* | 213 | 71 | 0.78 | 0.86 | 0.88 | 0.75 | 0.83 | 0.81 |
|  | *Cases* | 36 | 255 |  |  |  |  |  |  |
| **RUS** |  | *Controls* | *Cases* |  |  |  |  |  |  |
| **Reel Class** | *Controls* | 42 | 14 | 0.65 | 0.71 | 0.60 | 0.75 | 0.63 | 0.69 |
|  | *Cases* | 17 | 26 |  |  |  |  |  |  |

\* PPV: Positive Predictive Value, \*\* NPV: Negative Predictive Value

For MLP used with the clamping method, there is no difference between the methods used to get rid of the class imbalance in terms of classification performance (fig 7.)

**Figure 7.**
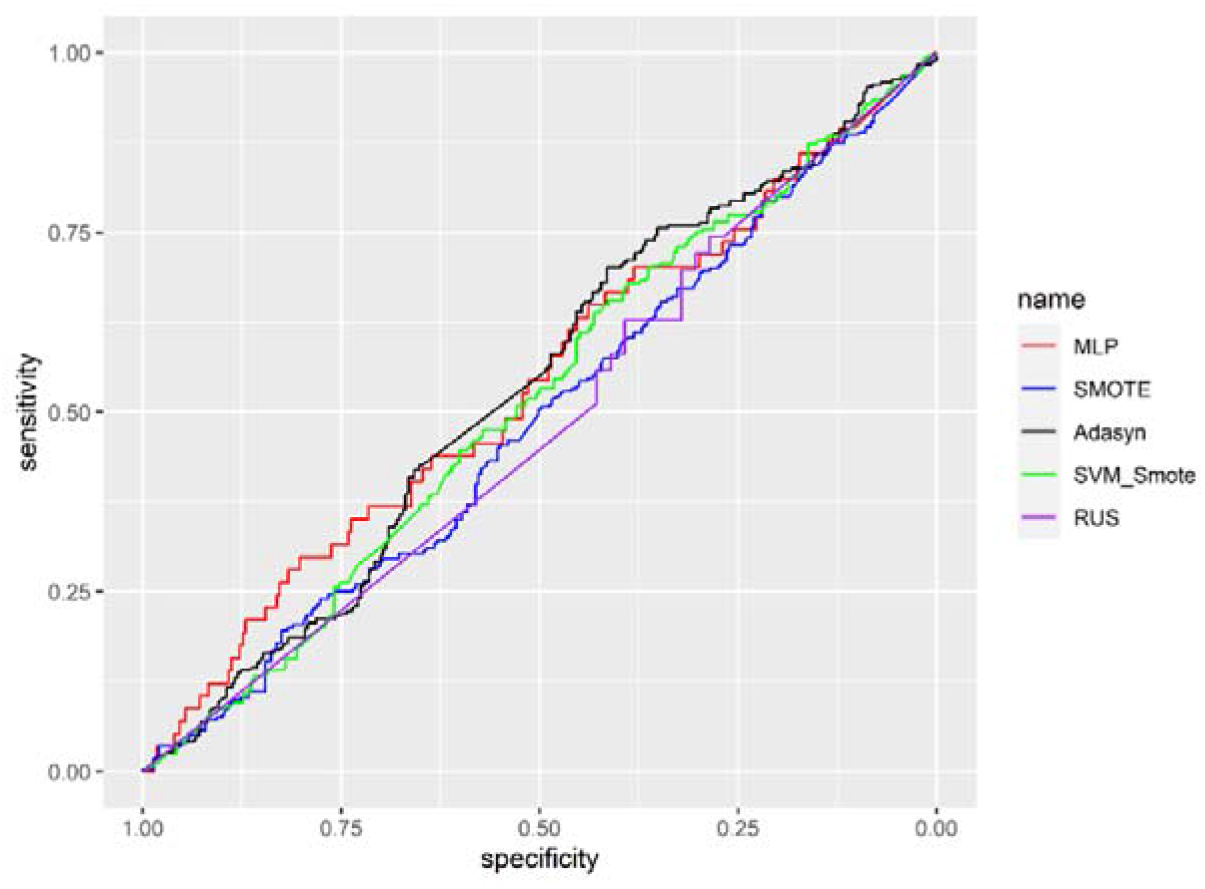
ROC Curve: Comparison of the performances of Multi-Layer Perceptron with Imbalanced Learning Methods with using Clumped SNPs

While high accuracy (89%) was obtained for data without class imbalance, low sensitivity rate (50%) and low F1 score (60%) were obtained due to class imbalance. In all the methods that eliminated the class imbalance, F1 score, and sensitivity were obtained at a higher rate than the original data, including RUS (Table 9).

**Table 9.**
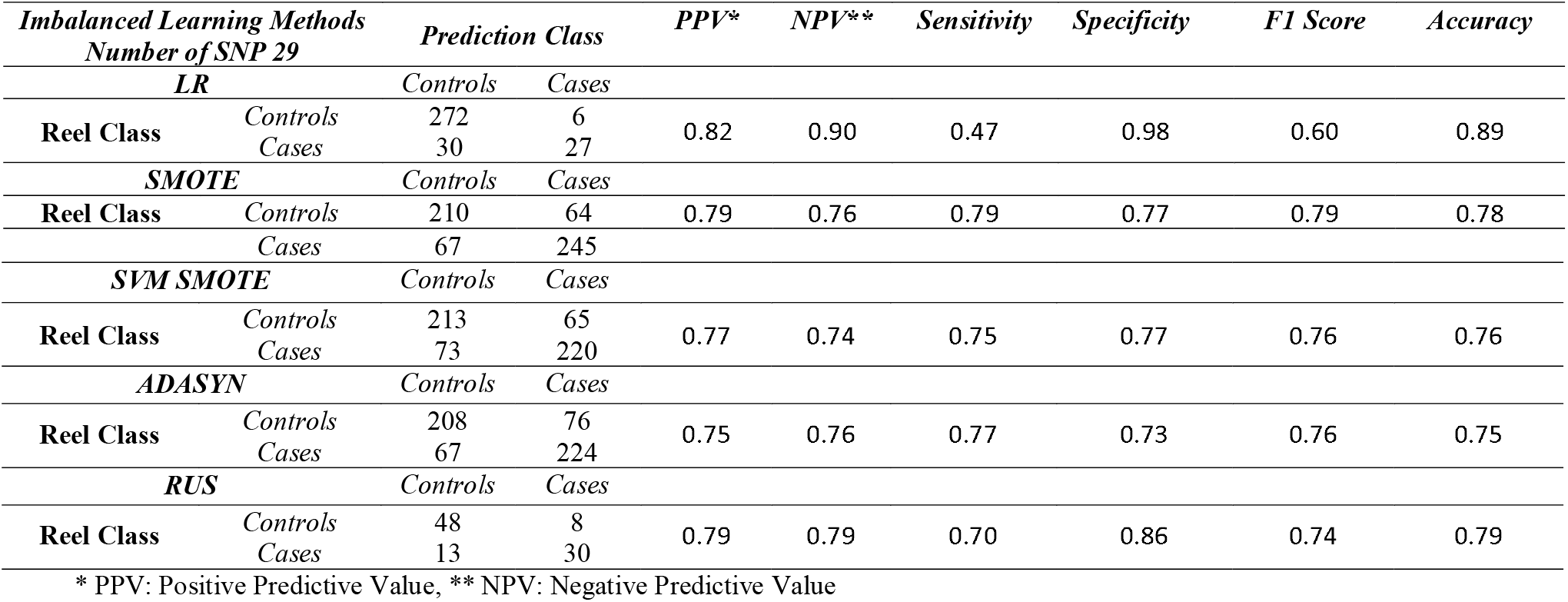
The Performances of Logistic Regression with Imbalanced Learning Methods with using Clumped SNPs.

All the methods used to eliminate class imbalance gave very close results for LR and LR is no different in terms of classification performance from the method used to eliminate any class imbalance. (fig. 8.)

**Figure 8.**
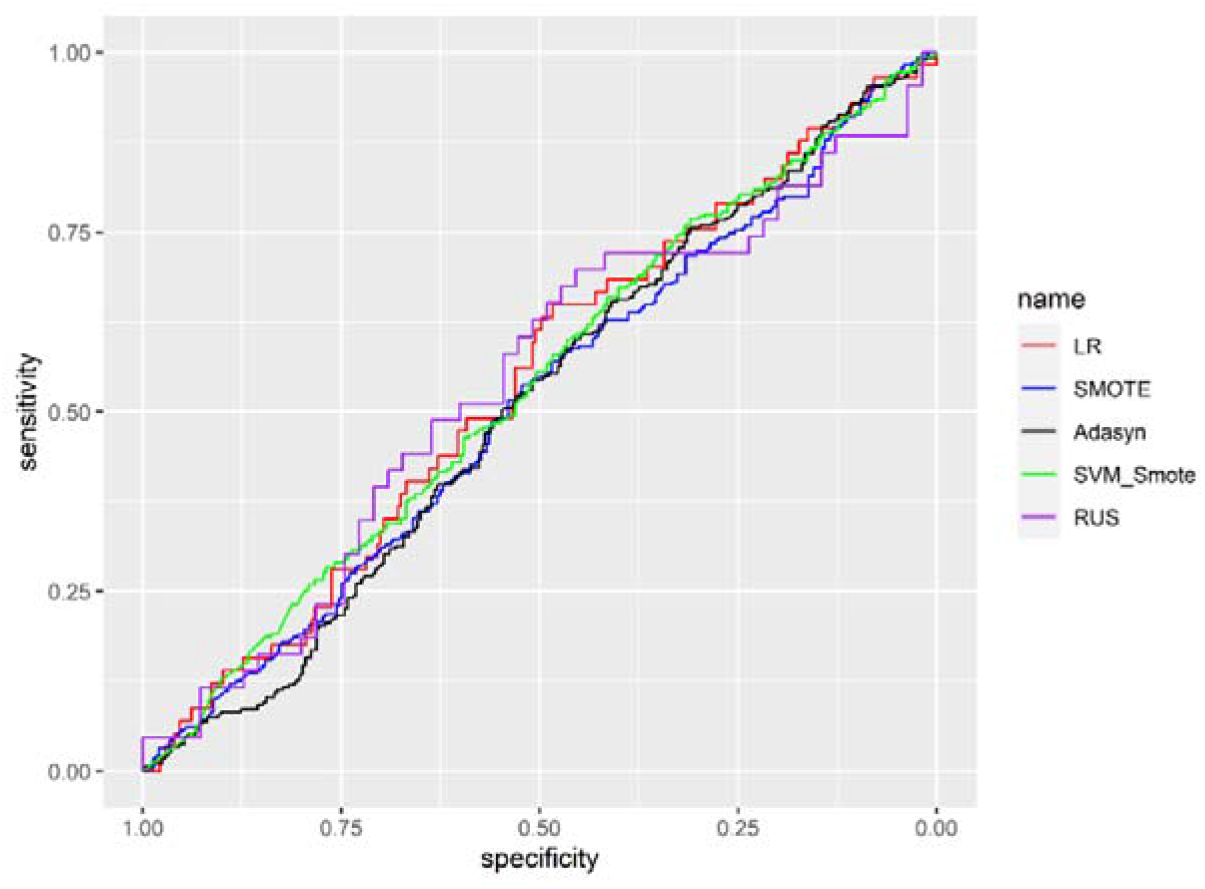
ROC Curve: Comparison of the performances of Logistic Regression with Imbalanced Learning Methods with using Clumped SNPs

### 3.3 Assessment of Machine Learning Results

As will be noted in all tables, accuracy is not the best indicator when evaluating ML models, whereas the F1 score is more informative. All models achieved “good” accuracy results and most methods achieved a positive predictive value and sensitivity value of 0.00. Zero precision algorithms cannot capture true positives. The methods used to resolve the class imbalance not only improved the performance of machine learning methods, but also increased the performance for the classical method, logistic regression. SMOTE applied to all SNPs has been shown to produce better results than clustering with SMOTE (Table 2–5). MLP was the best method overall, but all methods were performed similarly for each analysis comparison. Considering the clumping method, all machine learning methods used to get rid of class imbalance only increased the sensitivity (Table 6–9).

Since the methods we used to eliminate class imbalance in all methods gave close results and the method that gave the best results in general among them was SMOTE, when we compared the BMI correction using the SMOTE method with the BMI correction, it was seen that BMI had no effect on the classification success. As shown in Supplementary Tables 1-4, although BMI is a highly significant predictor of T2D, it has little effect on the predictive metrics evaluated across the methodology. Most interesting is the effect of BMI on the MLP algorithm when clustered SMOTE is applied as there is an increase in prediction accuracy. When we compare the MLP performance in Table 5 and Supplementary Table 3, we can see an increase in prediction accuracy of 0.05 and better classification of cases and controls.

### 3.4 Covariate Investigation

Obesity is a well-established risk factor for T2D. The mean and standard deviation (SD) of BMI in the controls (N=951) and cases (N=165) were 25.97±3,22 and 27.93±3,94 respectively. As expected, BMI is strongly associated with T2D (t=-6.035, p < 0.001). We repeated our classification with adjustment for BMI by including it as a variable in the model. The ML results including BMI are shown in the Supplementary Material. All the models both with and without clumping were not affected by the inclusion of BMI. Nevertheless, the same results were obtained for the models with SMOTE (Supplementary Tables 1-4).

## 4 DISCUSSION

The results when using SMOTE (resampling) for imbalanced classification are shown to be the most accurate. The clumping method is better suited in terms of computational speed but is dependent on LD pruning because correlated SNPs are less likely to be eliminated from the analysis. Only a small number of related SNPs were selected using clumping and then pruned by the chosen LD threshold.

Our findings suggest that using the SMOTE method with all the SNPs in a given dataset should be implemented in order to avoid over-fitting. Doing this enabled the use of the whole dataset for SNP pattern recognition compared to the clumping procedure, where many potentially false positive or true negative associated SNPs were not eliminated.

The specificity results in Table 4, highlight a concern with using ML methods. ML methods fall short when dealing with an imbalanced class of data for selecting controls. For this reason, methods such as clumping and SMOTE are important for achieving unbiased results (De Velasco Oriol, Vallejo, Estrada, Taméz Peña, & Initiative, 2019) (Privé, Vilhjálmsson, Aschard, & Blum, 2019), (Schubach, Re, Robinson, & Valentini, 2017). Because, when applying clumping, the SNPs that are lowly correlation with one another are included in the analysis. Applying SVM with clumping achieved very high classification metric results as shown in Table 6. As a final point, we recommend using SMOTE with any of the three ML models we discussed for GWAS data with imbalanced classes. This is because SMOTE seems unaffected by the imbalanced sample size and the volume of SNPs to be analysed. However further analysis is needed with a greater number of patients. Machine learning techniques are being applied to a variety of different data types and are growing in popularity in several industries. One key advantage over classical statistical methodology is that ML models do not require any assumptions. This helps in the search for patterns in large scale datasets produced in the field of genomics.

Different problems can be faced when using ML, such as class imbalance, computation time and memory usage. The imbalanced class problem causes underfitting, and we have shown that using SMOTE could be a solution (Li et al., 2016). Our results show that using SMOTE with RF can drastically improve prediction performance. On the other hand, clumping is beneficial for reducing computation time and memory usage with improved prediction performance over using ML methods without clumping. The clumping method is the best option for large datasets due to its stringent feature elimination criteria. The inflated accuracy results from all models may be related to the dimensionality of the data. Previous studies of a similar nature have shown comparable accuracy rates (T. Zheng et al., 2017), (Poplin et al., 2018), (Bao et al., 2021).

Among the methods used to eliminate the other class imbalance, SMOTE and ADASYN generally obtained close results. The RUS method, on the other hand, had poor results when used with all SNPs, because if examined, information loss may occur in randomly selected samples depending on the data size. The reason for the good results of using RUS with Clumping should be taken into account that the loss of information may be less because the data size is reduced (it has decreased to only 29 SNPs).

This paper makes recommendations and suggestions based on data similar to that of the ULSAM genetic and cohort data. Investigators should always be careful and mindful of the impact on their results when selecting and using ML models with imbalanced classes. Previous literature have shown that class imbalance can affect sensitivity or specificity (Afzal et al., 2013). ML is currently a useful method for the validation of classical GWAS approaches to identify SNPs associated with disease and for the prediction of disease status. As more and larger datasets become widely available, this will enable ML algorithms to predict disease status using SNP data more accurately in future cohorts. Leaving behind the need for stringent p-value thresholds and assumptions.

In this study, machine learning is not used for find out a new SNPs, it is used for diagnostics and precision medicine in individual. This study is a novel for GWAS data with Class-Imbalanced. Compared to other methods, we recommend using SMOTE with MLP.

## Supporting information

C:\Users\onuro\Desktop\Supplementary Files.docx

## REFERENCES

Afzal, Z., Schuemie, M. J., van Blijderveen, J. C., Sen, E. F., Sturkenboom, M. C., & Kors, J. A. (2013). Improving sensitivity of machine learning methods for automated case identification from free-text electronic medical records. BMC medical informatics and decision making, 13(1), 1–11.

Alpaydin, E. (2020). Introduction to machine learning: MIT press.

Bao, Z., Zhao, X., Li, J., Zhang, G., Wu, H., Ning, Y., … Yang, Z. (2021). Prediction of repeated-dose intravenous ketamine response in major depressive disorder using the GWAS-based machine learning approach. Journal of Psychiatric Research, 138, 284–290.

Bhowan, U., Johnston, M., Zhang, M., & Yao, X. (2012). Evolving diverse ensembles using genetic programming for classification with unbalanced data. IEEE Transactions on Evolutionary Computation, 17(3), 368–386.

Breiman, L. (2001). Random forests. Machine learning, 45(1), 5–32.

Chawla, N. V., Bowyer, K. W., Hall, L. O., & Kegelmeyer, W. P. (2002). SMOTE: synthetic minority over-sampling technique. Journal of artificial intelligence research, 16, 321–357.

Chen, X., & Ishwaran, H. (2012). Random forests for genomic data analysis. Genomics, 99(6), 323–329.

Chicco, D., & Jurman, G. (2020). The advantages of the Matthews correlation coefficient (MCC) over F1 score and accuracy in binary classification evaluation. BMC genomics, 21(1), 1–13.

Cosgun, E., Limdi, N. A., & Duarte, C. W. (2011). High-dimensional pharmacogenetic prediction of a continuous trait using machine learning techniques with application to warfarin dose prediction in African Americans. Bioinformatics, 27(10), 1384–1389.

Dai, X., Fu, G., Zhao, S., & Zeng, Y. (2021). Statistical Learning Methods Applicable to Genome-Wide Association Studies on Unbalanced Case-Control Disease Data. Genes, 12(5), 736.

De Velasco Oriol, J., Vallejo, E. E., Estrada, K., Taméz Peña, J. G., & Initiative, D. N. (2019). Benchmarking machine learning models for late-onset alzheimer’s disease prediction from genomic data. BMC bioinformatics, 20(1), 1–17.

Deng, F., Shen, L., Wang, H., & Zhang, L. (2020). Classify multicategory outcome in patients with lung adenocarcinoma using clinical, transcriptomic and clinico-transcriptomic data: machine learning versus multinomial models. American journal of cancer research, 10(12), 4624.

Draisma, H. H., Pool, R., Kobl, M., Jansen, R., Petersen, A.-K., Vaarhorst, A. A., … Esko, T. (2015). Genome-wide association study identifies novel genetic variants contributing to variation in blood metabolite levels. Nature communications, 6(1), 1–9.

Elmas, Ç., & Uygulamalari, Y. Z. (2007). Yapay Sinir Ağlari, Bulanik Mantik, Genetik Algoritmalar, 1. Basım, Ankara: Seçkin Yayincilik.

Fadista, J., Manning, A. K., Florez, J. C., & Groop, L. (2016). The (in) famous GWAS P-value threshold revisited and updated for low-frequency variants. European Journal of Human Genetics, 24(8), 1202–1205.

Fan, Y., & Tang, C. Y. (2013). Tuning parameter selection in high dimensional penalized likelihood. Journal of the Royal Statistical Society: Series B (Statistical Methodology), 75(3), 531–552.

Fergus, P., Montanez, C. C., Abdulaimma, B., Lisboa, P., Chalmers, C., & Pineles, B. (2018). Utilizing deep learning and genome wide association studies for epistatic-driven preterm birth classification in African-American women. IEEE/ACM transactions on computational biology and bioinformatics, 17(2), 668–678.

Furey, T. S., Cristianini, N., Duffy, N., Bednarski, D. W., Schummer, M., & Haussler, D. (2000). Support vector machine classification and validation of cancer tissue samples using microarray expression data. Bioinformatics, 16(10), 906–914.

Guyon, I., Weston, J., Barnhill, S., & Vapnik, V. (2002). Gene selection for cancer classification using support vector machines. Machine learning, 46(1), 389–422.

Han, J., Pei, J., & Kamber, M. (2011). Data mining: concepts and techniques: Elsevier.

He, K. Y., Ge, D., & He, M. M. (2017). Big data analytics for genomic medicine. International journal of molecular sciences, 18(2), 412.

Hu, F., & Li, H. (2013). A novel boundary oversampling algorithm based on neighborhood rough set model: NRSBoundary-SMOTE. Mathematical Problems in Engineering, 2013.

Japkowicz, N., & Stephen, S. (2002). The class imbalance problem: A systematic study. Intelligent data analysis, 6(5), 429–449.

Johnstone, I. M., & Titterington, D. M. (2009). Statistical challenges of high-dimensional data. In (Vol. 367, pp. 4237–4253): The Royal Society Publishing.

Kinreich, S., McCutcheon, V. V., Aliev, F., Meyers, J. L., Kamarajan, C., Pandey, A. K., … Pandey, G. (2021). Predicting alcohol use disorder remission: a longitudinal multimodal multi-featured machine learning approach. Translational psychiatry, 11(1), 1–10.

Korkmaz, S. (2020). Deep learning-based imbalanced data classification for drug discovery. Journal of Chemical Information and Modeling, 60(9), 4180–4190.

Lavesson, N., & Davidsson, P. (2006). Quantifying the impact of learning algorithm parameter tuning. Paper presented at the AAAI.

Li, J., Fong, S., Mohammed, S., Fiaidhi, J., Chen, Q., & Tan, Z. (2016). Solving the under-fitting problem for decision tree algorithms by incremental swarm optimization in rare-event healthcare classification. Journal of Medical Imaging and Health Informatics, 6(4), 1102–1110.

Lithell, H., Sundström, J., Ärnlöv, J., Björklund, K., Hänni, A., Hedman, A., … Reneland, R. (2000). Epidemiological and clinical studies on insulin resistance and diabetes. Upsala Journal of Medical Sciences, 105(2), 135–150.

Lusa, L. (2013). Improved shrunken centroid classifiers for high-dimensional class-imbalanced data. BMC bioinformatics, 14(1), 1–13.

Mammone, A., Turchi, M., & Cristianini, N. (2009). Support vector machines. Wiley Interdisciplinary Reviews: Computational Statistics, 1(3), 283–289.

Afzal, Z., Schuemie, M. J., van Blijderveen, J. C., Sen, E. F., Sturkenboom, M. C., & Kors, J. A. (2013). Improving sensitivity of machine learning methods for automated case identification from free-text electronic medical records. BMC Med Inform Decis Mak, 13(1), 30. doi:10.1186/1472-6947-13-30

Alhudhaif, A. (2021). A novel multi-class imbalanced EEG signals classification based on the adaptive synthetic sampling (ADASYN) approach. PeerJ Computer Science, 7, e523.

Alpaydin, E. (2020). Introduction to machine learning: MIT press.

Breiman, L. (2001). Random forests. Machine learning, 45(1), 5–32.

Elmas, Ç., & Uygulamalaπ, Y. Z. (2007). Yapay Sinir Ağlaπ, Bulamk Mantik, Genetik Algoritmalar, 1. Basım, Ankara: Seçkin Yaymcilik.

Han, J., Pei, J., & Kamber, M. (2011). Data mining: concepts and techniques: Elsevier.

He, H., Bai, Y., Garcia, E. A., & Li, S. (2008). ADASYN: Adaptive synthetic sampling approach for imbalanced learning. Paper presented at the 2008 IEEE international joint conference on neural networks (IEEE world congress on computational intelligence).

Mieth, B., Kloft, M., Rodríguez, J. A., Sonnenburg, S., Vobruba, R., Morcillo-Suárez, C., … Dickhaus, T. (2016). Combining multiple hypothesis testing with machine learning increases the statistical power of genome-wide association studies. Scientific reports, 6(1), 1–14.

Ng, K. L. S., & Mishra, S. K. (2007). De novo SVM classification of precursor microRNAs from genomic pseudo hairpins using global and intrinsic folding measures. Bioinformatics, 23(11), 1321–1330.

Nitze, I., Schulthess, U., & Asche, H. (2012). Comparison of machine learning algorithms random forest, artificial neural network and support vector machine to maximum likelihood for supervised crop type classification. Proceedings of the 4th GEOBIA, Rio de Janeiro, Brazil, 79, 3540.

Nordhausen, K. (2009). The elements of statistical learning: data mining, inference, and prediction, by trevor hastie, robert tibshirani, jerome friedman. In: Wiley Online Library.

Pal, M. (2005). Random forest classifier for remote sensing classification. International journal of remote sensing, 26(1), 217–222.

Pal, S. K., & Mitra, S. (1992). Multilayer perceptron, fuzzy sets, classifiaction.

Pedregosa, F., Varoquaux, G., Gramfort, A., Michel, V., Thirion, B., Grisel, O., … Dubourg, V. (2011). Scikit-learn: Machine learning in Python, the Journal of machine Learning research, 12, 2825–2830.

Pirooznia, M., Fayaz Seifuddin, J. J., Mahon, P. B., Potash, J. B., Zandi, P. P., & Consortium, B. G. S. (2012). Data mining approaches for genome-wide association of mood disorders. Psychiatric genetics, 22(2), 55.

Poplin, R., Chang, P.-C., Alexander, D., Schwartz, S., Colthurst, T., Ku, A., … Afshar, P. T. (2018). Creating a universal SNP and small indel variant caller with deep neural networks. BioRxiv, 092890.

Privé, F., Vilhjálmsson, B. J., Aschard, H., & Blum, M. G. (2019). Making the most of clumping and thresholding for polygenic scores. The American journal of human genetics, 105(6), 1213–1221.

Purcell, S., Neale, B., Todd-Brown, K., Thomas, L., Ferreira, M. A., Bender, D., … Daly, M. J. (2007). PLINK: a tool set for whole-genome association and population-based linkage analyses. The American journal of human genetics, 81(3), 559–575.

Razavi-Far, R., Farajzadeh-Zanajni, M., Wang, B., Saif, M., & Chakrabarti, S. (2019). Imputation-based ensemble techniques for class imbalance learning. IEEE Transactions on Knowledge and Data Engineering, 33(5), 1988–2001.

Schubach, M., Re, M., Robinson, P. N., & Valentini, G. (2017). Imbalance-aware machine learning for predicting rare and common disease-associated non-coding variants. Scientific reports, 7(1), 1–12.

Seo, J.-H., & Kim, Y.-H. (2018). Machine-learning approach to optimize smote ratio in class imbalance dataset for intrusion detection. Computational intelligence and neuroscience, 2018.

Shi, H., Medway, C., Brown, K., Kalsheker, N., & Morgan, K. (2011). Using Fisher’s method with PLINK ‘LD clumped’output to compare SNP effects across Genome-wide Association Study (GWAS) datasets. International journal of molecular epidemiology and genetics, 2(1), 30.

Shrivastava, S., Jeyanthi, P. M., & Singh, S. (2020). Failure prediction of Indian Banks using SMOTE, Lasso regression, bagging and boosting. Cogent Economics & Finance, 8(1), 1729569.

Staley, J. R., Jones, E., Kaptoge, S., Butterworth, A. S., Sweeting, M. J., Wood, A. M., & Howson, J. M. (2017). A comparison of Cox and logistic regression for use in genome-wide association studies of cohort and case-cohort design. European Journal of Human Genetics, 25(1), 854–862.

Statnikov, A., Aliferis, C. F., Tsamardinos, L., Hardin, D., & Levy, S. (2005). A comprehensive evaluation of multicategory classification methods for microarray gene expression cancer diagnosis. Bioinformatics, 21(5), 631–643.

Strobl, C., & Zeileis, A. (2008). Danger: High power!-exploring the statistical properties of a test for random forest variable importance.

Szymczak, S., Biernacka, J. M., Cordell, H. J., González-Recio, O., König, I. R., Zhang, H., & Sun, Y. V. (2009). Machine learning in genome-wide association studies. Genetic epidemiology, 33(S1), S51–S57.

Tang, Y., Zhang, Y.-Q., Chawla, N. V., & Krasser, S. (2008). SVMs modeling for highly imbalanced classification. IEEE Transactions on Systems, Man, and Cybernetics, Part B (Cybernetics), 39(1), 281–288.

Turhan, S., Özkan, Y., Yürekli, B. S., Suner, A., & Doğu, E. (2020). Sinif Dengesizliği Varliğinda Hastahk Tanisi için Kolektif Öğrenme Yöntemlerinin Karşilaştirilmasi: Diyabet Tanisi Örneği. Turkiye Klinikleri Journal of Biostatistics, 12(1).

Van Rossum, G. (2007). Python Programming language. Paper presented at the USENIX annual technical conference.

Wakefield, J. (2009). Bayes factors for genome-wide association studies: comparison with P-values. Genetic Epidemiology: The Official Publication of the International Genetic Epidemiology Society, 33(1), 79–86.

Wang, H.-Y. (2008). Combination approach of SMOTE and biased-SVM for imbalanced datasets. Paper presented at the 2008 IEEE International Joint Conference on Neural Networks (IEEE World Congress on Computational Intelligence).

Wang, Q., Luo, Z., Huang, J., Feng, Y., & Liu, Z. (2017). A novel ensemble method for imbalanced data learning: bagging of extrapolation-SMOTE SVM. Computational intelligence and neuroscience, 2017.

Zheng, T., Xie, W., Xu, L., He, X., Zhang, Y., You, M., … Chen, Y. (2017). A machine learning-based framework to identify type 2 diabetes through electronic health records. International journal of medical informatics, 97,120-127.

Zheng, Z., Cai, Y., & Li, Y. (2015). Oversampling method for imbalanced classification. Computing and Informatics, 34(5), 1017–1037.

Zhou, W., Nielsen, J. B., Fritsche, L. G., Dey, R., Gabrielsen, M. E., Wolford, B. N., … Gifford, A. (2018). Efficiently controlling for case-control imbalance and sample relatedness in large-scale genetic association studies. Nature genetics, 50(9), 1335–1341.

Zuech, R., Hancock, J., & Khoshgoftaar, T. M. (2021). Detecting web attacks using random undersampling and ensemble learners. Journal of Big Data, 8(1), 1–20.

Pal, S. K., & Mitra, S. (1992). Multilayer perceptron, fuzzy sets, classifiaction.

Pedregosa, F., Varoquaux, G., Gramfort, A., Michel, V., Thirion, B., Grisel, O., … Dubourg, V. (2011). Scikit-learn: Machine learning in Python. the Journal of machine Learning research, 12, 2825–2830.

Statnikov, A., Aliferis, C. F., Tsamardinos, I., Hardin, D., & Levy, S. (2005). A comprehensive evaluation of multicategory classification methods for microarray gene expression cancer diagnosis. Bioinformatics, 21(5), 631–643.

Strobl, C., & Zeileis, A. (2008). Danger: High power!–exploring the statistical properties of a test for random forest variable importance.

Szymczak, S., Biernacka, J. M., Cordell, H. J., González□Recio, O., König, I. R., Zhang, H., & Sun, Y. V. (2009). Machine learning in genome□wide association studies. Genetic epidemiology, 33(S1), S51–S57.

Turhan, S., Özkan, Y., Yürekli, B. S., Suner, A., & Doğu, E. (2020). Sinif Dengesizliği Varliğinda Hastalik Tanisi için Kolektif Öğrenme Yöntemlerinin Karşilaştirilmasi: Diyabet Tanisi Öπieġi. Turkiye Klinikleri Journal of Biostatistics, 12(1).

Thomas, M., Sakoda, L. C., Hoffmeister, M., Rosenthal, E. A., Lee, J. K., van Duijnhoven, F. J., … de la Chapelle, A. (2020). Genome-wide modeling of polygenic risk score in colorectal cancer risk. The American journal of human genetics, 107(3), 432–444.

Zheng, T., Xie, W., Xu, L., He, X., Zhang, Y., You, M., … Chen, Y. (2017). A machine learning-based framework to identify type 2 diabetes through electronic health records. International journal of medical informatics, 97, 120–127.

