## Supplementary material for "The Use of Class Imbalanced Learning Methods on ULSAM Data to Predict the Case-Control Status in Genome-Wide Association Studies": C:\Users\onuro\Desktop\Supplementary Files.docx

Tuning Parameters Table

| Tuning Parameters Table | | | | |
| --- | --- | --- | --- | --- |
| Methods | Criteria | Parameters | Optimization Range | Best Results |
| Support  Vector  Machine | Radial Kernel | C | [0:100] * | C=1 |
| Random  Forest | Gini,  Bootstrap | Max_depth, n_estimators | max_depth= [10:100]*  n_estimators= [0:2000] ** | max_depth=30, n_estimators=1000 |
| Multilayer Perceptron | Activation  Relu | α, ꞵ, epsilon, batch, epochs | batchs= [10:120] *  iter= [0:100] * | α=1e-05, beta1=0.9, beta2=0.999, epsilon=1e-08  hidden_layer_sizes=(20, 4) |

* Tried every 10-step interval, ** Tried every 100-step interval.

BMI tables

***Table 1. The Performances of Machine Learning Methods with SMOTE***

| Classification Methods  Number of SNP 399,935 | | Prediction Class | | PPV* | NPV** | Sensitivity | Specificity | F1 Score | Accuracy |
| --- | --- | --- | --- | --- | --- | --- | --- | --- | --- |
| *Support Vector Machine* | | *Controls* | *Cases* |  | | | | |  |
| Actual Class | *Controls* | 275 | 0 | 1.00 | 0.84 | 0.82 | 1.00 | 0.90 | 0.91 |
|  | *Cases* | 51 | 245 |  |  |  |  |  |  |
| *Random Forest* | | *Controls* | *Cases* |  | | | | |  |
| Actual Class | *Controls* | 275 | 0 | 1.00 | 0.84 | 0.82 | 1.00 | 0.90 | 0.91 |
|  | *Cases* | 51 | 245 |  |  |  |  |  |  |
| *Multi-Layer Perceptron* | | *Controls* | *Cases* |  | | | | |  |
| Actual Class | *Controls* | 274 | 1 | 0.99 | 0.90 | 0.90 | 0.99 | 0.94 | 0.94 |
|  | *Cases* | 28 | 268 |  |  |  |  |  |  |

* PPV: Positive Predictive Value, ** NPV: Negative Predictive Value

***Table 2. The Performances of Machine Learning Methods without using SMOTE***

| *Classification Methods*  *Number of SNP 399,935* | | *Prediction Class* | | *PPV** | *NPV*** | *Sensitivity* | *Specificity* | *F1 Score* | *Accuracy* |
| --- | --- | --- | --- | --- | --- | --- | --- | --- | --- |
| *Support Vector Machine* | | *Controls* | *Cases* |  | | | | |  |
| Actual  Class | *Controls* | 278 | 0 | 0.00 | 0.82 | 0.00 | 1.00 | 0.00 | 0.82 |
|  | *Cases* | 57 | 0 |  |  |  |  |  |  |
| *Random Forest* | | *Controls* | *Case* | “ | | | | |  |
| Actual  Class | *Controls* | 278 | 0 | 0.00 | 0.82 | 0.00 | 1.00 | 0.00 | 0.82 |
|  | *Cases* | 57 | 0 |  |  |  |  |  |  |
| *Multi-Layer Perceptron* | | *Controls* | *Cases* |  | | | | |  |
| Actual  Class | *Controls* | 278 | 0 | 1.00 | 0.83 | 0.01 | 1.00 | 0.03 | 0.83 |
|  | *Cases* | 56 | 1 |  |  |  |  |  |  |

* PPV: Positive Predictive Value, ** NPV: Negative Predictive Value

**Clumped**

***Table 3. The Performances of Machine Learning Methods without using SMOTE***

| *Classification Methods*  *Number of SNP 29* | | Prediction Class | | *PPV** | *NPV*** | *Sensitivity* | *Specificity* | *F1 Score* | *Accuracy* |
| --- | --- | --- | --- | --- | --- | --- | --- | --- | --- |
| *Support Vector Machine* | | *Controls* | *Cases* |  | | | | |  |
| Actual  Class | *Controls* | 279 | 0 | 1.00 | 0.83 | 0.05 | 1.00 | 0.10 | 0.83 |
|  | *Cases* | 54 | 3 |  |  |  |  |  |  |
| *Random Forest* | | *Controls* | *Cases* |  | | | | |  |
| Actual  Class | *Controls* | 279 | 0 | 0.00 | 0.83 | 0.00 | 1.00 | 0.00 | 0.83 |
|  | *Cases* | 57 | 0 |  |  |  |  |  |  |
| *Multi-Layer Perceptron* | | *Controls* | *Cases* |  | | | | |  |
| Actual  Class | *Controls* | 270 | 9 | 0.71 | 0.88 | 0.40 | 0.96 | 0.51 | 0.87 |
|  | *Cases* | 34 | 23 |  |  |  |  |  |  |

* PPV: Positive Predictive Value, ** NPV: Negative Predictive Value

***Table 4. The Performances of Machine Learning Methods with using SMOTE***

| *Classification Methods*  *Number of SNP 29* | | Prediction Class | | *PPV** | *NPV*** | *Sensitivity* | *Specificity* | *F1 Score* | *Accuracy* |
| --- | --- | --- | --- | --- | --- | --- | --- | --- | --- |
| *Support Vector Machine* | | *Controls* | *Cases* |  | | | | |  |
| Actual  Class | *Controls* | 216 | 63 | 0.93 | 0.77 | 0.81 | 0.91 | 0.86 | 0.85 |
|  | *Cases* | 20 | 273 |  |  |  |  |  |  |
| *Random Forest* | | *Controls* | *Cases* |  | | | | |  |
| Actual  Class | *Controls* | 247 | 32 | 0.91 | 0.88 | 0.89 | 0.90 | 0.90 | 0.89 |
|  | *Cases* | 26 | 267 |  |  |  |  |  |  |
| *Multi-Layer Perceptron* | | *Controls* | *Cases* |  | | | | |  |
| Actual  Class | *Controls* | 200 | 79 | 0.76 | 0.84 | 0.87 | 0.71 | 0.81 | 0.79 |
|  | *Cases* | 38 | 255 |  |  |  |  |  |  |

* PPV: Positive Predictive Value, ** NPV: Negative Predictive Value

**Table 5. Significant SNPs using Clumping**

| CHR | F | SNP | BP | P | TOTAL | NSIG | S05 | S01 | S001 | S0001 | SP2 |
| --- | --- | --- | --- | --- | --- | --- | --- | --- | --- | --- | --- |
| 14 | 1 | rs11850812 | 55183330 | 3.79e-07 | 0 | 0 | 0 | 0 | 0 | 0 | NONE |
| 16 | 1 | rs12597362 | 13753349 | 5.65e-07 | 0 | 0 | 0 | 0 | 0 | 0 | NONE |
| 5 | 1 | rs16869618 | 90536573 | 6.75e-07 | 2 | 0 | 0 | 0 | 1 | 1 | rs16869554(1),rs12520306(1) |
| 1 | 1 | rs12094313 | 81278056 | 6.79e-07 | 1 | 0 | 0 | 0 | 0 | 1 | rs11163137(1) |
| 10 | 1 | rs11591913 | 80673648 | 2.61e-06 | 0 | 0 | 0 | 0 | 0 | 0 | NONE |
| 14 | 1 | rs854344 | 25340520 | 5.17e-06 | 0 | 0 | 0 | 0 | 0 | 0 | NONE |
| 12 | 1 | rs11063482 | 5159686 | 1.2e-05 | 0 | 0 | 0 | 0 | 0 | 0 | NONE |
| 18 | 1 | rs11083124 | 22987297 | 1.24e-05 | 2 | 0 | 2 | 0 | 0 | 0 | NONE |
| 10 | 1 | rs2505956 | 27185286 | 1.24e-05 | 2 | 0 | 1 | 0 | 1 | 0 | rs2182296(1) |
| 16 | 1 | rs7203539 | 79494293 | 1.29e-05 | 1 | 0 | 0 | 0 | 1 | 0 | rs2549516(1) |
| 12 | 1 | rs2446994 | 49238024 | 1.34e-05 | 1 | 0 | 0 | 1 | 0 | 0 | rs2453481(1) |
| 2 | 1 | rs717027 | 1.67E+08 | 1.37e-05 | 2 | 0 | 0 | 0 | 1 | 1 | rs13413443(1),rs9677856(1) |
| 3 | 1 | rs6767775 | 84157105 | 1.65e-05 | 4 | 0 | 0 | 2 | 2 | 0 | rs6548932(1),rs4605562(1),rs6767473(1),rs6548940(1) |
| 12 | 1 | rs17124060 | 61080039 | 1.74e-05 | 0 | 0 | 0 | 0 | 0 | 0 | NONE |
| 11 | 1 | - | 61570073 | 2.17e-05 | 0 | 0 | 0 | 0 | 0 | 0 | NONE |
| 11 | 1 | rs2155466 | 1.21E+08 | 2.17e-05 | 1 | 0 | 0 | 0 | 1 | 0 | rs4936580(1) |
| 11 | 1 | rs4362115 | 38031231 | 2.33e-05 | 6 | 0 | 0 | 3 | 3 | 0 | rs1679407(1),rs11034567(1),rs12296091(1),rs11034572(1),rs821011(1),rs7947901(1) |
| 12 | 1 | rs6489601 | 5140530 | 2.46e-05 | 2 | 0 | 0 | 0 | 1 | 1 | rs12099807(1),rs6489602(1) |
| 4 | 1 | rs2866787 | 1.07E+08 | 2.57e-05 | 0 | 0 | 0 | 0 | 0 | 0 | NONE |
| 6 | 1 | rs9341424 | 74170860 | 3.04e-05 | 9 | 0 | 1 | 5 | 1 | 2 | rs535902(1),rs674708(1),rs4708060(1),rs364734(1),rs685491(1),rs687210(1),rs472294(1),rs4708072(1) |
| 5 | 1 | rs1437118 | 94301042 | 3.05e-05 | 3 | 0 | 0 | 1 | 2 | 0 | rs6890714(1),rs10043438(1),rs13180001(1) |
| 20 | 1 | rs3746755 | 61578610 | 3.57e-05 | 2 | 0 | 0 | 1 | 0 | 1 | rs1883847(1),rs1052826(1) |
| 6 | 1 | rs6924865 | 26521353 | 3.61e-05 | 46 | 0 | 1 | 5 | 19 | 21 | rs2073526(1),rs1796520(1),rs1796521(1),rs4320356(1),rs1624440(1),rs9295689(1),rs1977198(1),rs7763910(1),rs1407045(1),rs6926629(1),rs3736781(1),rs1056667(1),rs9393729(1),rs1535276(1),rs9295695(1),rs10946835(1),rs2393670(1),rs9467782(1),rs4871(1),rs9986382(1),rs12526680(1),rs2224380(1),rs9461271(1),rs10484442(1),rs767471(1),rs6456733(1),rs1078679(1),rs6925895(1),rs1321482(1),rs1570061(1),rs6918854(1),rs6925087(1),rs4368798(1),rs3800303(1),rs9467810(1),rs2451738(1),rs2451731(1),rs2451741(1),rs1027204(1),rs2172007(1),rs2504599(1),rs2504571(1),rs2504600(1),rs2504565(1),rs2498399(1) |
| 10 | 1 | rs946517 | 80529202 | 3.79e-05 | 5 | 2 | 0 | 1 | 2 | 0 | rs7910759(1),rs7904670(1),rs11002673(1) |
| 18 | 1 | rs11662423 | 74180990 | 4.45e-05 | 1 | 0 | 0 | 1 | 0 | 0 | rs661859(1) |
| 18 | 1 | rs2698158 | 68604686 | 4.91e-05 | 2 | 0 | 0 | 2 | 0 | 0 | rs1395855(1),rs1605451(1) |
| 12 | 1 | rs2247919 | 1.03E+08 | 5.04e-05 | 1 | 0 | 0 | 0 | 0 | 1 | rs1722402(1) |
| 9 | 1 | rs13287994 | 84370124 | 5.05e-05 | 0 | 0 | 0 | 0 | 0 | 0 | NONE |
| 8 | 1 | rs7006025 | 59355053 | 5.08e-05 | 8 | 0 | 0 | 2 | 5 | 1 | rs10957052(1),rs11776617(1),rs4541896(1),rs16923491(1),rs6471722(1),  rs10957056(1),rs8192877(1),rs11786580(1) |
| 1 | 1 | rs12123975 | 1.52E+08 | 6.1e-05 | 4 | 0 | 0 | 0 | 4 | 0 | rs3007671(1),rs2999547(1),rs12134184(1),rs17646946(1) |
| 18 | 1 | rs7234673 | 68036279 | 6.42e-05 | 0 | 0 | 0 | 0 | 0 | 0 | NONE |
| 13 | 1 | rs11840821 | 42162235 | 6.7e-05 | 3 | 0 | 0 | 3 | 0 | 0 | rs4299049(1),rs7989696(1),rs7999090(1) |
| 4 | 1 | rs6847657 | 1.02E+08 | 6.87e-05 | 4 | 0 | 0 | 1 | 3 | 0 | rs10008442(1),rs13130138(1),rs6840909(1),rs17826594(1) |
| 2 | 1 | rs7586417 | 1.43E+08 | 7.31e-05 | 3 | 0 | 0 | 1 | 0 | 2 | rs1449479(1),rs1449480(1),rs10193058(1) |
| 15 | 1 | rs8042430 | 1.02E+08 | 7.55e-05 | 0 | 0 | 0 | 0 | 0 | 0 | NONE |
| 4 | 1 | rs3946 | 367927 | 7.62e-05 | 9 | 0 | 2 | 2 | 4 | 1 | rs11946825(1),rs6825544(1),rs11725919(1),rs11734441(1),rs7695712(1),  rs6810900(1),rs17721347(1) |
| 7 | 1 | rs1581532 | 67791971 | 7.67e-05 | 1 | 0 | 0 | 1 | 0 | 0 | rs10266576(1) |
| 22 | 1 | rs5992749 | 18076523 | 7.8e-05 | 0 | 0 | 0 | 0 | 0 | 0 | NONE |
| 1 | 1 | rs11205983 | 53102133 | 7.8e-05 | 0 | 0 | 0 | 0 | 0 | 0 | NONE |
| 2 | 1 | rs16826971 | 1.48E+08 | 7.9e-05 | 0 | 0 | 0 | 0 | 0 | 0 | NONE |
| 10 | 1 | rs6585063 | 1.13E+08 | 8.38e-05 | 3 | 0 | 0 | 2 | 1 | 0 | rs3885682(1),rs10787335(1),rs7903217(1) |
| 14 | 1 | rs1824343 | 98920773 | 8.89e-05 | 1 | 0 | 0 | 1 | 0 | 0 | rs17097078(1) |
| 17 | 1 | rs2239921 | 43170907 | 8.98e-05 | 2 | 0 | 0 | 1 | 1 | 0 | chr17:40526433(1),chr17:40542083(1) |
| 1 | 1 | rs16837837 | 2.39E+08 | 9.00E-05 | 2 | 0 | 0 | 2 | 0 | 0 | rs17641617(1),rs17567797(1) |
| 5 | 1 | rs7729395 | 1.02E+08 | 9.14e-05 | 0 | 0 | 0 | 0 | 0 | 0 | NONE |
| 12 | 1 | rs1544502 | 2438810 | 9.17e-05 | 7 | 0 | 0 | 5 | 2 | 0 | rs2108570(1),rs2239041(1),rs3819526(1),rs3819536(1),rs2283301(1),  rs2283302(1),rs2238070(1) |
| 4 | 1 | rs11944122 | 442021 | 9.17e-05 | 0 | 0 | 0 | 0 | 0 | 0 | NONE |
| 3 | 1 | rs6767011 | 36504281 | 9.34e-05 | 0 | 0 | 0 | 0 | 0 | 0 | NONE |
| 16 | 1 | rs902553 | 86387524 | 9.98e-05 | 1 | 0 | 0 | 0 | 0 | 1 | rs1687677(1) |

CHR : Chromosome code, F	: Results fileset code (1,2,...), SNP : SNP identifier, BP	: Physical position of SNP (base-pairs),
TOTAL: Total number of other SNPs in clump (i.e. passing --clump-kb and --clump-r2 thresholds), NSIG : Number of clumped SNPs that are not significant ( p > 0.05 ), S05 : Number of clumped SNPs 0.01 < p < 0.05, S01 : Number of clumped SNPs 0.001 < p < 0.01, S001 : Number of clumped SNPs 0.0001 < p < 0.001, S0001: Number of clumped SNPs p < 0.0001, SP2 : List of SNPs names (and fileset code) clumped and significant at --clump-p2
